# The Sleep-Mood Connection: Temperament Modulates Neuroinflammation, Clock Genes, and Dopaminergic Receptors Expression in Rats

**DOI:** 10.1101/2025.06.09.658634

**Authors:** Camila Nayane Carvalho Lima, Francisco Eliclécio Rodrigues da Silva, Michele Albuquerque Jales de Carvalho, Jose Eduardo de Carvalho Lima, Adriano José Maia Chaves-Filho, Gabriel R. Fries, Manoel A. Sobreira-Neto, Deiziane Viana da Silva Costa, Marta Maria França Fonteles, Danielle S. Macêdo

## Abstract

Temperament may influence mood disorder risk under sleep disturbances. Using high- and low-exploratory rats as a temperament model, we examined the effects of paradoxical sleep deprivation (PSD) on mood behaviors, neuroinflammation, oxidative stress, clock genes, and dopamine receptor expression. Eighty male Wistar rats were screened for exploratory activity, while 20 were classified as high (HE) and 21 as low (LE) exploratory based on open-field performance. HE and LE rats were subjected to PSD or remained undisturbed (controls). Behavioral assessments included impulsivity, risk-taking, anhedonia, and forced swim test (FST) performance. Neurobiological analyses measured hippocampal lipid peroxidation, pro-inflammatory markers (IL-6, TLR4), gene expression of *Clock*, *Tph2, Tdo2,* and protein expression of D1, and D2, which regulate, respectively, circadian rhythms, serotonin synthesis, tryptophan metabolism, and dopaminergic signaling—key pathways implicated in mood disorders. HE rats showed increased impulsivity, risk-taking, and exploratory behaviors, resembling mania, while LE rats exhibited avoidance. PSD intensified these traits, heightening risk-taking in HE rats and inducing depressive-like behaviors in LE rats, including anhedonia and FST immobility linked to reduced Tph2 expression. Both groups had post-PSD working memory deficits. Neurobiologically, HE+PSD rats showed increased lipid peroxidation, neuroinflammation (IL-6, TLR4), and upregulation of *Clock* and DRD1 genes, while LE+PSD rats had elevated hippocampal *Tdo2* expression. Temperament shapes PSD-induced behavioral and neurobiological effects. HE rats showed mania-like traits, while LE rats exhibited depressive behaviors. PSD impaired working memory in both. Oxidative stress, inflammation, and gene expression changes highlight distinct vulnerabilities and potential therapeutic targets for mood disorders.

## 1. Introduction

Temperament refers to innate personality tendencies, including emotional reactivity, affective expressiveness, and sensitivity (Merikangas et al., 1998). It is biologically driven and stable throughout life and shapes exploratory behavior in humans and mammals. Rodent models offer key insights into harm avoidance and novelty seeking (Steimer and Driscoll, 2005). Harm avoidance involves fearfulness and timidity(Cloninger et al., 1993a), while novelty seeking involves curiosity and exploration(Kazlauckas et al., 2005).

Understanding individual differences in psychiatric vulnerability is challenging. Given temperament’s role in mental disorders, exploring its neurobiological mechanisms through rodent behavior in novel environments is essential (Whittle et al., 2006). For this purpose, high- (HE) and low-exploratory (LE) rats serve as useful models to simulate novelty seeking and harm avoidance, respectively (Görisch and Schwarting, 2006; Pawlak and Schwarting, 2002; Thiel et al., 1999).

In a previous study, HE and LE Wistar rats exposed to repetitive paradoxical sleep deprivation (PSD) and unpredictable stress (US) exhibited distinct behavioral and immune changes. HE rats showed anxiolytic-like behavior, while both groups displayed memory impairments. Immune dysregulation, marked by Th1/Th2 imbalance, contributed to neuroinflammation and oxidative stress. Lithium treatment partially mitigated these effects (Lima et al., 2019).

Circadian rhythms and clock genes are implicated in mood disorders, regulating physiological processes that influence mood (Jones, 2001). Disruptions in these rhythms can cause mood fluctuations seen in bipolar disorder (BD) and depression. These fluctuations often coincide with sleep-wake cycle and other circadian function alterations (Jagannath et al., 2013; Rumble et al., 2015a). Normalizing sleep/wake cycles and social interactions is crucial for mood stabilization (Rumble et al., 2015b). Mania, a fundamental feature of BD, involves hyperactivity, reduced anxiety, and increased risk-taking (Phillips and Kupfer, 2013). Manipulation of *Clock* (Roybal et al., 2007a) or related molecules *Per1* and *Per2* (Abarca et al., 2002), specifically within the mesolimbic dopamine system, produced animals with an ‘‘HE-like’’ behavioral phenotype.

The hippocampus, crucial for learning and memory, expresses core circadian clock genes like *CLOCK, Bmal1, Per, and Cry*. These genes regulate local oscillations affecting synaptic plasticity, memory consolidation, and mood regulation (A.Kondratova et al., 2010; Wang et al., 2019). The circadian clock operates through a biochemical feedback loop in mammals, where *Bmal1* and *Clock* activate *Per* and *Cry* transcription. Per and Cry proteins then inhibit *Clock*: *Bmal1* activity, creating a cycle essential for generating circadian rhythms (Landgraf et al., 2014; Mcclung, 2011). These genes also regulate immune responses and, in the hippocampus, dopamine synthesis and receptor activity (Wang and Li, 2021). Disruptions in circadian genes can alter dopaminergic signaling, closely linked to mood disorders like depression and BD (Jiang et al., 2019). Investigating biological clocks may reveal links between exploratory behavior in rats, environmental stressors, and mood disorders.

This study hypothesized that HE and LE rats subjected to PSD would exhibit distinct behavioral phenotypes, resembling BD and depression, respectively. These behavioral differences were proposed to be driven by alterations in clock gene expression (*Clock*), neuroinflammatory markers [interleukin-6 (IL-6), toll-like receptor 4 (TLR-4)], oxidative stress (lipid peroxidation), serotonin synthesis (tryptophan hydroxylase 2; *Tph2* expression), tryptophan metabolism (tryptophan 2,3-dioxygenase; *Tdo2* expression), and dopamine receptor (DR) 1 and 2 protein expression. The primary objective of this study was to evaluate behavioral changes in HE and LE rats with and without PSD exposure. The secondary objective was to investigate the underlying neurobiological mechanisms contributing to these behavioral alterations.

## 2. Methods and Materials

2.1. **Animals**

Male periadolescent Wistar rats (50–60 g) from the Universidade Federal do Ceará Animal House were housed in groups of five under controlled conditions (22 ± 1°C, 60 ± 5% humidity, 12-h light/dark cycle, lights on at 7:00 a.m.) with ad libitum access to food and water. Blinded raters conducted behavioral tests between 8:00–14:00 h. All procedures complied with Brazilian regulations and NIH guidelines, were approved by the Ethics Committee (protocol 125/14), and aimed to minimize suffering and animal use.

### 2.2. Behavioral analyses of HE- and LE- rats

2.2.1. **Screening for locomotor response to novelty**

A psychogenic selection method was used to classify periadolescent rats (postnatal day – PND - 30) into high (HE) and low (LE) exploratory groups based on a 10-minute open field test (25th and 75th percentiles). Exploratory profiles remained stable in adulthood (PND 60). Of 80 screened rats, 20 HE (≥75th percentile, >67.25% rearing) and 20 LE (≤25th percentile, <32% rearing) were selected, while 40 median rats were excluded. HE rats showed higher exploratory behavior than LE rats [(HE: 80.60 ± 9.99%, N=20); (LE: 21.90 ± 7.55%, N=21); t=21.29, df=39, p<0.0001]. Reassessment at PND 60 confirmed stable exploratory profiles (t=8.25, p<0.0001) (Supplementary Table 1, Figure 3B and C). All tests were conducted between ZT2 (8 a.m.) and ZT5 (11 a.m.).

#### 2.2.2. Experimental Protocol

After HE and LE classification at PND30, on PND40-49 rats were randomly assigned to PSD or control conditions. PSD followed a 10-day protocol (alternate days) in groups of five per cage (41 cm × 34 cm × 16.5 cm) with 12 small platforms (3 cm diameter) over shallow water, inducing awakening upon muscle atonia. Control rats were housed similarly but without water. Food and water were available ad libitum (Gessa et al., 1995).

Rats were divided into four groups (n = 8/group): HE-control, HE+PSD, LE-control, and LE+PSD. Behavioral assessments included risk-taking (cat odor), locomotion (open-field test - OFT), anxiety (elevated plus maze - EPM), depression-like behavior (forced swim test - FST), anhedonia (sucrose preference - SP), working memory (Y-maze), and declarative memory (novel object recognition).

After euthanasia, blood and brain tissue were analyzed for uric acid, glutathione (GSH), lipid peroxidation, nitrite levels, and IL-1β, IL-4, and IL-6 immunoassays. Gene expression of *Clock, Bmal1, Cry1, Cry2, Per1, Per2, Per3,* and protein levels of D1 and D2 receptors were also assessed.

### 2.3 Behavioral parameters analyzed

The aversive exposure to cat odor test measured risk-taking behavior by recording cloth contact and sheltering behavior when rats were exposed to either a predator odor or neutral cloth (Zangrossi and File, 1992).

The OFT assessed locomotion and exploration in a 50 × 50 cm arena, analyzing center avoidance, central-to-peripheral locomotion ratio, immobility, quadrant transitions, and rearing. The EPM test measured anxiety by quantifying the percentage of open-arm entries and time spent in open arms of a maze elevated 50 cm above the floor (Pellow et al., 1985).

The FST performed in a 60 cm-high water-filled cylinder, evaluated depressive-like behavior based on immobility duration across two sessions (10 min initial exposure, 5 min re-exposure) (Muscat and Willner, 1992; Porsolt et al., 1978). The SPT (Scheggi et al., 2018) assessed anhedonia by measuring sucrose intake relative to total liquid consumption after 20 h of food and water deprivation.

Cognitive function was evaluated using the Y-maze test, which measured spatial working memory through spontaneous alternation in a three-armed maze (SARTER et al., 1988), and the novel object recognition (NOR) test (Izquierdo et al., 2006), where rats explored two identical objects, followed by the replacement of one with a novel object after 1 or 5 hours, with the recognition index calculated as the proportion of time spent exploring the novel object. All tests were recorded using video tracking software, and appropriate cleaning procedures were applied between trials to prevent olfactory cues.

Figure 1 illustrates the experimental timeline and behavioral paradigms employed in this study.

**Figure 1.**
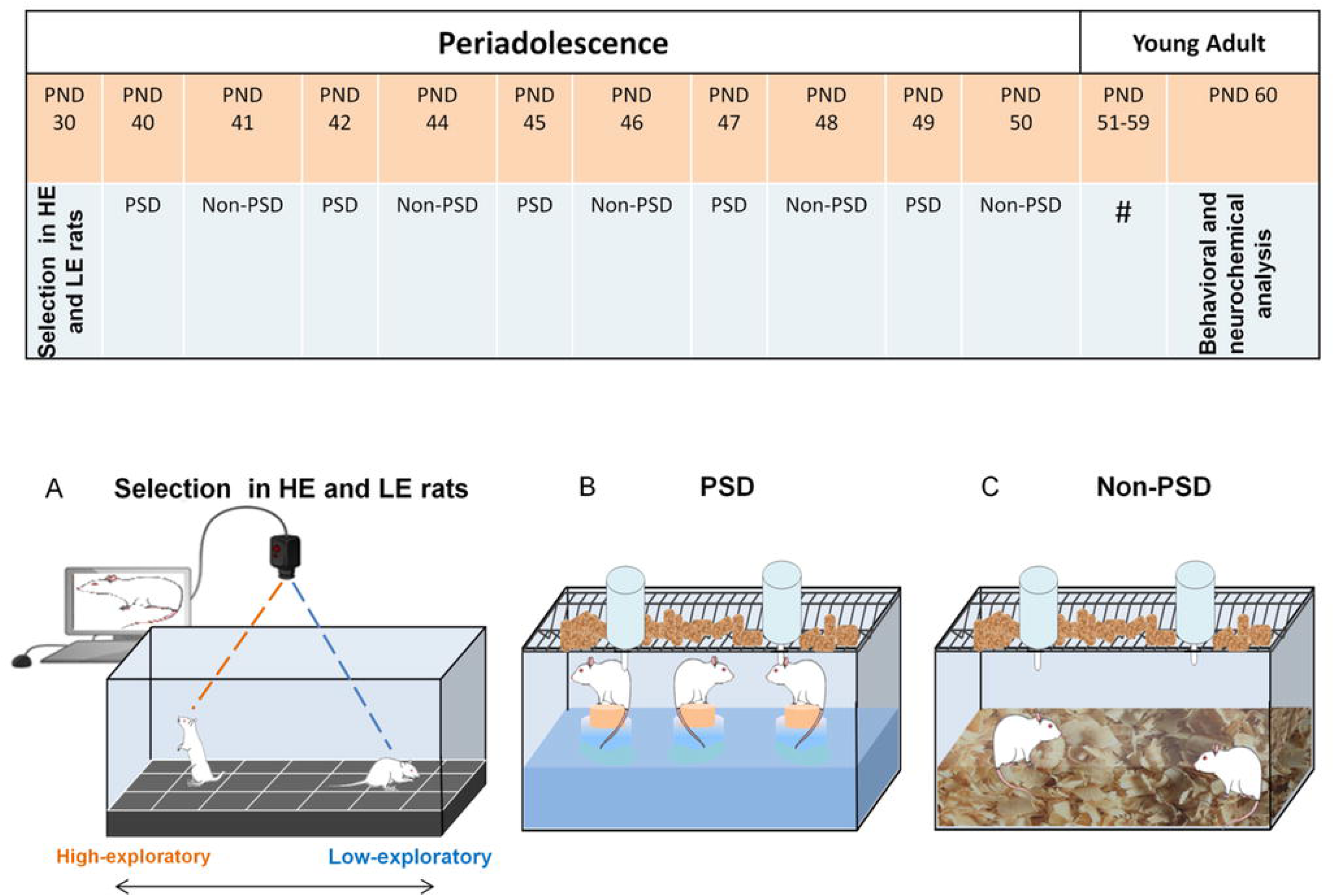
**Experimental timeline and behavioral paradigms for High-exploratory (HE) and Low-exploratory (LE) rats during periodolescence and young adulthood** A psychogenic selection was employed to study temperament in rodents by dividing them into extremes of exploratory activity. Periadolescent rats on the 30th postnatal day (PND30) were selected based on their performance in a 10-minute open field test session. The rats were divided into two groups: HE (high-exploratory) and LE (low-exploratory) rats, categorized by their 25th and 75th percentile vertical exploratory activity levels. From PND40-49, these animals were randomly selected to be exposed to PSD (paradoxical sleep deprivation) or not, undergoing a ten-day protocol with PSD on alternate days. Behavioral and neurochemical analyses were performed on PND60.

### 2.4 Determination of oxidative and inflammatory parameters

#### 2.4.1 Uric Acid Levels and Oxidative Stress Determinations

Uric acid levels were routinely monitored in the rats’ serum samples using a commercial kit (Randox, San Diego, California).

Brain reduced glutathione (GSH) levels were measured at 412 nm using Ellman’s reagent (DTNB) to detect free thiol groups and expressed as ng GSH/g wet tissue (Uchiyama and Mihara, 1978). Lipid peroxidation was assessed via thiobarbituric acid reactive substances (TBARS) (Reilly and Aust, 1999). TBARS were quantified by absorbance at 532 nm, using a malondialdehyde (MDA) standard curve, and expressed as μmol MDA/mg tissue. Nitrite levels in brain homogenates, indirectly indicating nitric oxide (NO) production, were assessed using the Griess reaction (Radenovic and Selakovic, 2005; Tannenbaum et al., 1978) Absorbance (550 nm) was measured via microplate reader. Nitrite levels were quantified using a NaNO₂ standard curve (0.75–100 μM) and expressed as μmol/g protein (Tsikas, 2007).

#### 2.4.2 Immunoassay for IL-1β, IL-4, IL-6

The brain areas were homogenized in PBS buffer (1:8 w/v) with a protease inhibitor (EMD Biosciences) and centrifuged (10,000 rpm, 5 min). Cytokine concentrations in 50 μL samples were measured using ELISA (R&D Systems, Minneapolis, MN, USA) per the manufacturer’s protocol and expressed in pg/g tissue.

### 2.5 Gene expression analyses

Total RNA was isolated (PROMEGA) and quantified (NanoDrop, 260/280 >1.8). cDNA was synthesized from 1 µg RNA using the High-Capacity cDNA Reverse Transcription Kit (Applied Biosystems).

*m*RNA expression was analyzed by quantitative PCR (qPCR) using SYBR Green and primers from Applied Biosystems (sequences in Supplementary Table 2). 50 ng cDNA was added to *Clock, Bmal1, Cry1, Cry2, Per1, Per2, Per3, or Gapdh* assays in a 20 µL reaction. The thermocycler protocol was 50°C (2 min), 95°C (10 min), followed by 40 cycles of 95°C (15 s) and 60°C (60 s). Gene expression was normalized to *Gapdh* and analyzed using the ΔΔCt method.

### 2.6 Western Blotting

Hippocampi were homogenized in RIPA buffer (pH 7.6) with protease inhibitors, centrifuged (17 min, 4°C, 13,000 rpm), and supernatants collected. Protein concentrations were determined by the Bradford method. For Western blot, 20 µg protein (Laemmli buffer, 95°C, 5 min) was separated on 10% SDS-PAGE, transferred to PVDF membranes, and blocked with 5% BSA (1h, RT). Membranes were incubated overnight with primary antibodies (D1, D2, α-tubulin), followed by secondary antibodies. Bands were detected using ECL (Bio-Rad ChemiDoc), and densitometry was analyzed with ImageJ (NIH).

### 2.7. Data analysis

HE and LE rats were classified based on rearing frequency in the novel open field, with the upper half assigned to HE and the lower half to LE (see Section 2.4.1). These groups were used for behavioral and neurochemical analyses.

Student’s t-test was applied to continuous, symmetrically distributed data, while two-way ANOVA (factors: Exploratory Phenotype [HE, LE] and PSD Exposure [PSD, non-PSD]) was used for behavioral tests. RT-PCR results were analyzed using one-way ANOVA with the Tukey post hoc test. Data are presented as mean ± SEM, except for EP results (mean ± SD) for graphical clarity.

## 3. RESULTS

### 3.1 Uric Acid Levels Correlate with Impulsivity in HE and LE Rats

HE-non-PSD and HE+PSD rats showed higher uric acid levels than their LE counterparts (HE-non-PSD vs. LE-non-PSD, P = 0.0090; HE+PSD vs. LE+PSD, P < 0.0001; F₁,₁₉ = 7.048, P = 0.015, Figure 2A).

**Figure 2.**
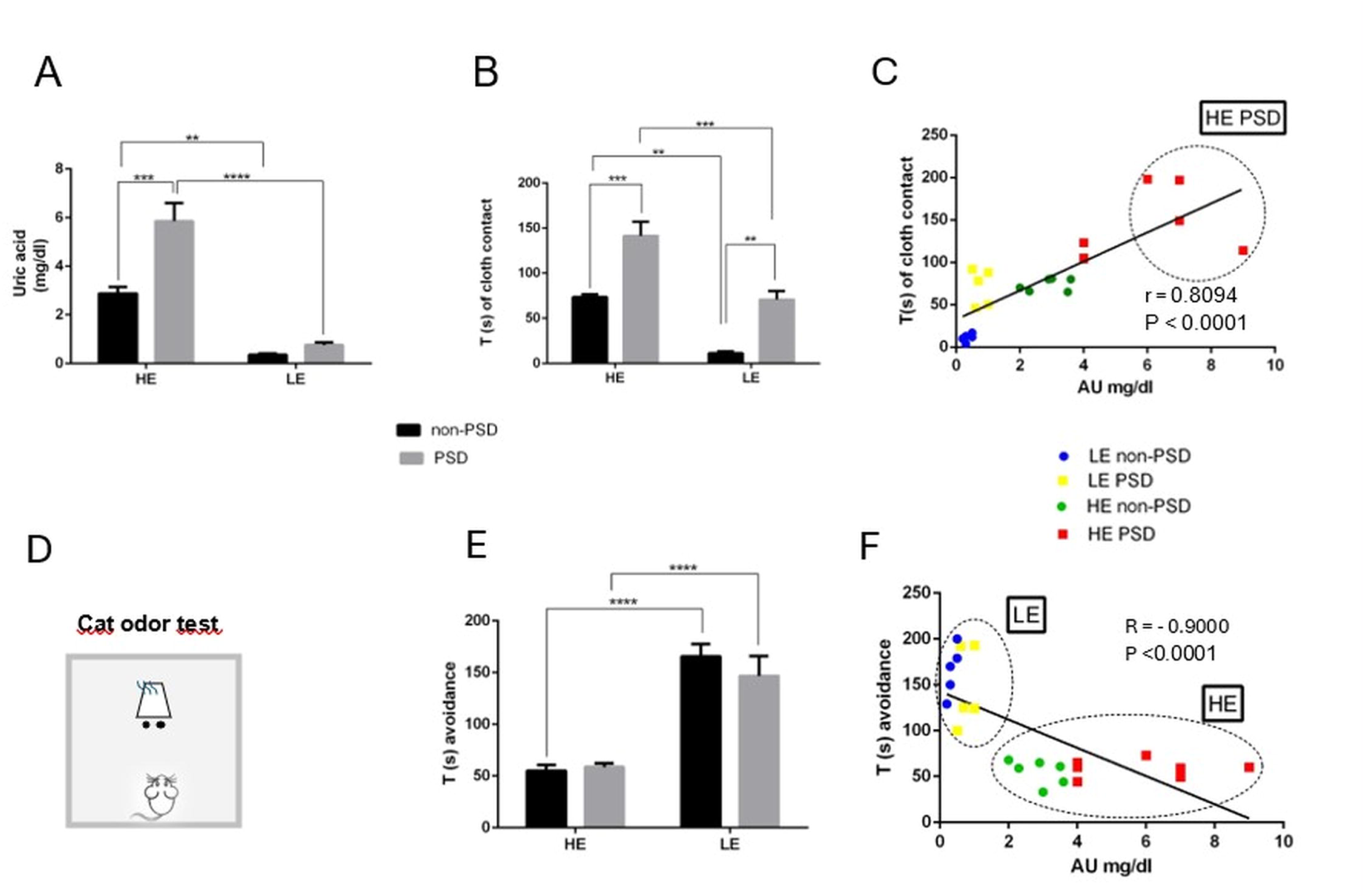
**Behavioral and biochemical analysis of High-exploratory (HE) and Low-exploratory (LE) rats: Effects of paradoxical sleep deprivation (PSD) on urate levels, cloth contact, and avoidance behavior** HE- and HE-PSD rats showed increased exploration and novelty-seeking behaviors in the cat odor-exposure test. (A) HE-PSD rats displayed a significant increase in serum uric acid (UA) levels compared to LE-PSD rats. (B) (C) In the cat odor-exposure test, HE- and HE-PSD rats had significantly spent more time in contact with the cloth than their respective LE- and LE-PSD rats, which was positively correlated with levels of uric acid. (D) Cat odor testing apparatus. (E) (F) Duration of avoidance was significantly higher for LE compared to HE and significantly higher for LE-PSD compared to HE-PSD, which was negatively correlated with levels of uric acid. Each bar represents the mean ± S.E.M of 8– 10 animals/group. Data were analyzed by one-way ANOVA, with Tukey as a post hoc test. **P < 0.01, ***P < 0.001, ***P < 0.0001.

**Figure 3.**
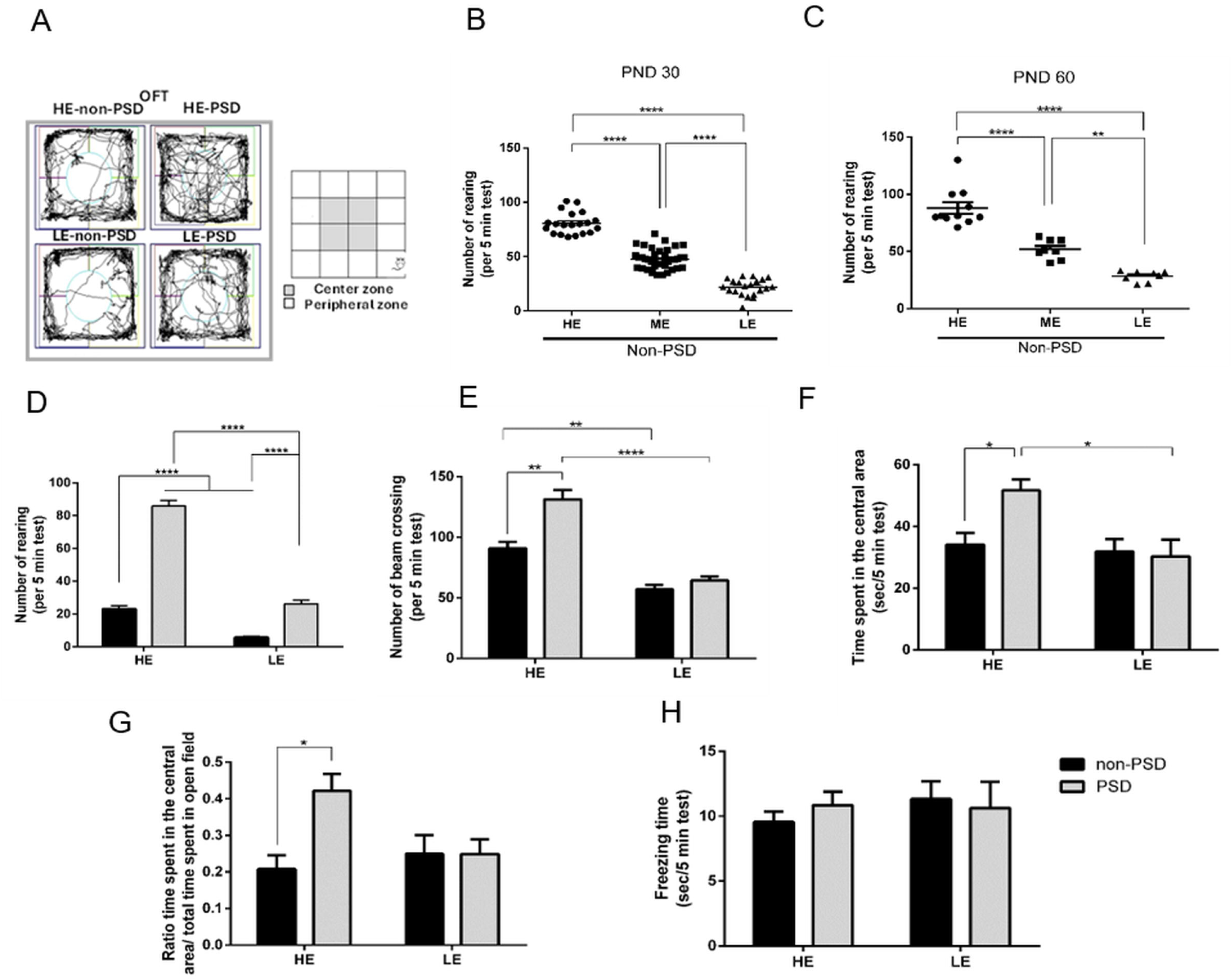
**Exploratory and locomotor behavior in High-exploratory (HE) and Low-exploratory (LE) Rats: Impact of paradoxical sleep deprivation (PSD) on activity and anxiety responses** (A) Representation of the animals’ paths in the central and peripheral zones during the behavioral tests. The groups are LE-non-PSD (low-exploratory, no PSD), LE-PSD (low-exploratory, with PSD), HE-non-PSD (high-exploratory, no PSD), and HE-PSD (high-exploratory, with PSD). (B and C) Number of runs per minute in the open field test at postnatal days (PND) 30 (B) and 60 (C), comparing the four groups. (D) Number of rearing movements per 5 minutes in the tests. (E) Number of beam crossings (locomotor activity) per 5 minutes. (F) Time spent in the central zone of the open field (seconds per minute), indicating anxiety behavior. (G) Ratio of time spent in the central zone versus the peripheral zone during the open field test. (H) Freezing time (in seconds per 5 minutes), indicating fear response. For all analyses, *p* < 0.05, \**p* < 0.01, ****p < 0.0001. Comparisons between the groups were made using appropriate statistical tests. The results show that paradoxical sleep deprivation induces significant alterations in locomotor activity and anxiety and fear-related behaviors, with differences observed based on exposure levels and the presence or absence of PSD.

HE rats spent more time in contact with the cat-odor cloth than LE rats (HE-non-PSD vs. LE-non-PSD, P = 0.0036; HE+PSD vs. LE+PSD, P = 0.0008, Figure 2B), which correlated positively with uric acid levels (R = 0.8094, P < 0.0001, Figure 2C). Conversely, avoidance duration was higher in LE rats (LE-non-PSD vs. HE-non-PSD, P < 0.0001; LE+PSD vs. HE+PSD, P < 0.0001, Figure 2E) and negatively correlated with uric acid (R = -0.9000, P < 0.0001, Figure 2F). A schematic representation of the cat-odor test is shown in Figure 2D.

### 3.2 PSD Differentially Modulates Exploratory Behavior in HE and LE Rats

We investigated whether PSD alters exploratory behavior in HE and LE rats (Figure 3A, for representative trajectories). We observed that the number of rearings remained stable in HE, ME, and LE rats on PND30 and 60 (Figure 3B and C). Notably, HE-non-PSD rats exhibited higher rearing (P < 0.0001, Figure 3D) and crossings (P = 0.0022, Figure 3E) than LE-non-PSD rats. PSD increased rearing frequency in both groups compared to non-PSD rats (F₁,₂₆ = 79.74, P < 0.0001, Figure 3D). However, only HE rats exhibited increased crossings (F₁,₁₈ = 8.568, P = 0.0090, Figure 3E), time spent in the central area (F₁,₁₈ = 5.019, P = 0.0379, Figure 3F), and central/peripheral time ratio (F₁,₁₈ = 5.661, P = 0.0286, Figure 3G) following PSD. Freezing time remained unchanged across groups (Figure 3H).

### 3.3 PSD Differentially Affects Anxiety, Depression, and Anhedonia in HE and LE Rats

We assessed PSD-induced anxiety-like behavior in HE and LE rats using the EPM test (Figure 4A). Both “exploratory phenotype” and “PSD exposure” influenced open-arm entries (F₁,₃₀ = 8.252, P = 0.0074) and time spent in open arms (F₁,₃₀ = 10.47, P = 0.0030). In this regard, HE-non-PSD rats exhibited higher risk-taking behavior, with more open-arm entries (P = 0.0017, Figure 4B) and longer open-arm time (P = 0.0157, Figure 4C) than LE-non-PSD rats. PSD increased these behaviors in HE rats (open-arm entries, P < 0.0001; open-arm time, P < 0.0001). In contrast, LE rats showed no PSD-induced changes (Figures 4B-C). For closed-arm entries (Figure 4D), HE-non-PSD rats displayed higher entries than LE-non-PSD rats (P < 0.0001). Additionally, HE+PSD rats spent a greater percentage of time in closed arms compared to LE-non-PSD rats (P < 0.0001).

**Figure 4.**
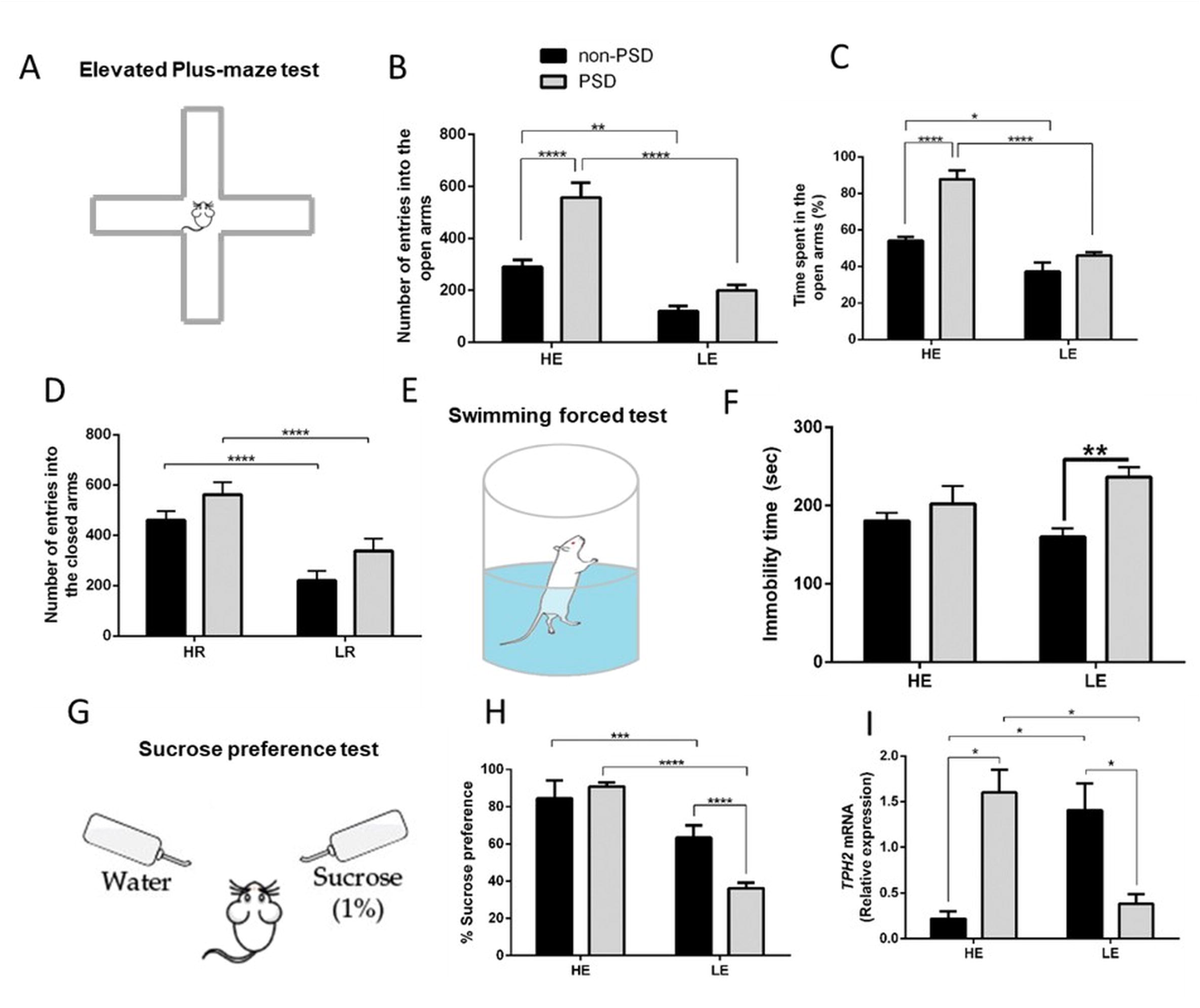
**Anxiety, depressive-like behavior, and reward sensitivity in High-exploratory (HE) and Low-exploratory (LE) rats: Impact of PSD on behavioral and molecular markers** (A) Elevated Plus-Maze (EPM) testing apparatus (B) Number of entries into the open arms of the EPM test, showing increased exploration in PSD groups, especially in High-exploratory (HE) compared to Low-exploratory (LE) groups. (C) Percentage of time spent in the open arms, indicating anxiety levels; a significant reduction in the LE-PSD group is observed compared to non-PSD controls. (D) Number of entries into the closed arms of the EPM test, indicating anxiety behaviors, with higher entries observed in the non-PSD groups. (E) Swimming Forced Test (SFT) apparatus (F) Time spent in immobility during the forced swim test, reflecting despair-like behavior, with significant differences between PSD and non-PSD groups in both HE and LE groups. (G) Sucrose Preference Test (SPT) apparatus (H) Percentage preference for sucrose solution in the sucrose preference test, indicating anhedonia, with reduced sucrose preference observed in PSD groups. (I) mRNA expression levels of *Tph2* (tryptophan hydroxylase 2) in the brain, showing significant differences in gene expression across the different groups. *p < 0.05, **p < 0.01, ***p < 0.001, ****p < 0.0001. Data were analyzed using appropriate statistical tests. The results indicate that PSD induces significant changes in anxiety-like behaviors, despair-like responses, and anhedonia, with variations observed based on exposure levels and PSD status.

Depression- and anhedonia-like behaviors induced by PSD in HE and LE rats were assessed using the forced swim test (FST, Figure 4E) and sucrose preference test (SPT, Figure 4G). A significant interaction between “Exploratory Phenotype” and “PSD exposure” was observed in SPT (F₁,₁₈ = 37.32, P < 0.0001), while FST was affected by PSD independently of phenotype (F₁,₂₄ = 10.96, P = 0.0029).

LE+PSD rats showed increased immobility time compared to LE-non-PSD rats (P = 0.0051, Figure 4F), whereas HE rats were unaffected by PSD. Sucrose preference decreased in LE+PSD rats compared to LE-non-PSD (P = 0.0001), which was already lower than in HE-non-PSD rats (P < 0.0001, Figure 4H). Similarly, LE+PSD rats had lower sucrose preference than HE+PSD rats (P < 0.0001), indicating greater vulnerability of LE rats to depression and anhedonia.

This was supported by *Tph2* mRNA levels, which decreased in LE+PSD rats compared to LE-non-PSD (P = 0.0454, Figure 4I), whereas HE+PSD rats exhibited increased *Tph2* expression relative to HE-non-PSD (P = 0.0193).

### 3.4 PSD Impairs Memory and Alters Oxidative Stress in HE and LE Rats

To assess the effects of PSD on working and declarative memory in HE- and LE-rats, we conducted Y-maze (Figure 5A) and NOR tests (Figure 5C). PSD impaired working memory in both phenotypes [F(1,27) = 8.241, P = 0.0079], but not declarative memory [F(1,19) = 4.190, P = 0.0548]. However, declarative memory was independently influenced by “Exploratory phenotype” [F(1,19) = 24.39, P < 0.0001] and “PSD exposure” [F(1,19) = 130.9, P < 0.0001].

**Figure 5.**
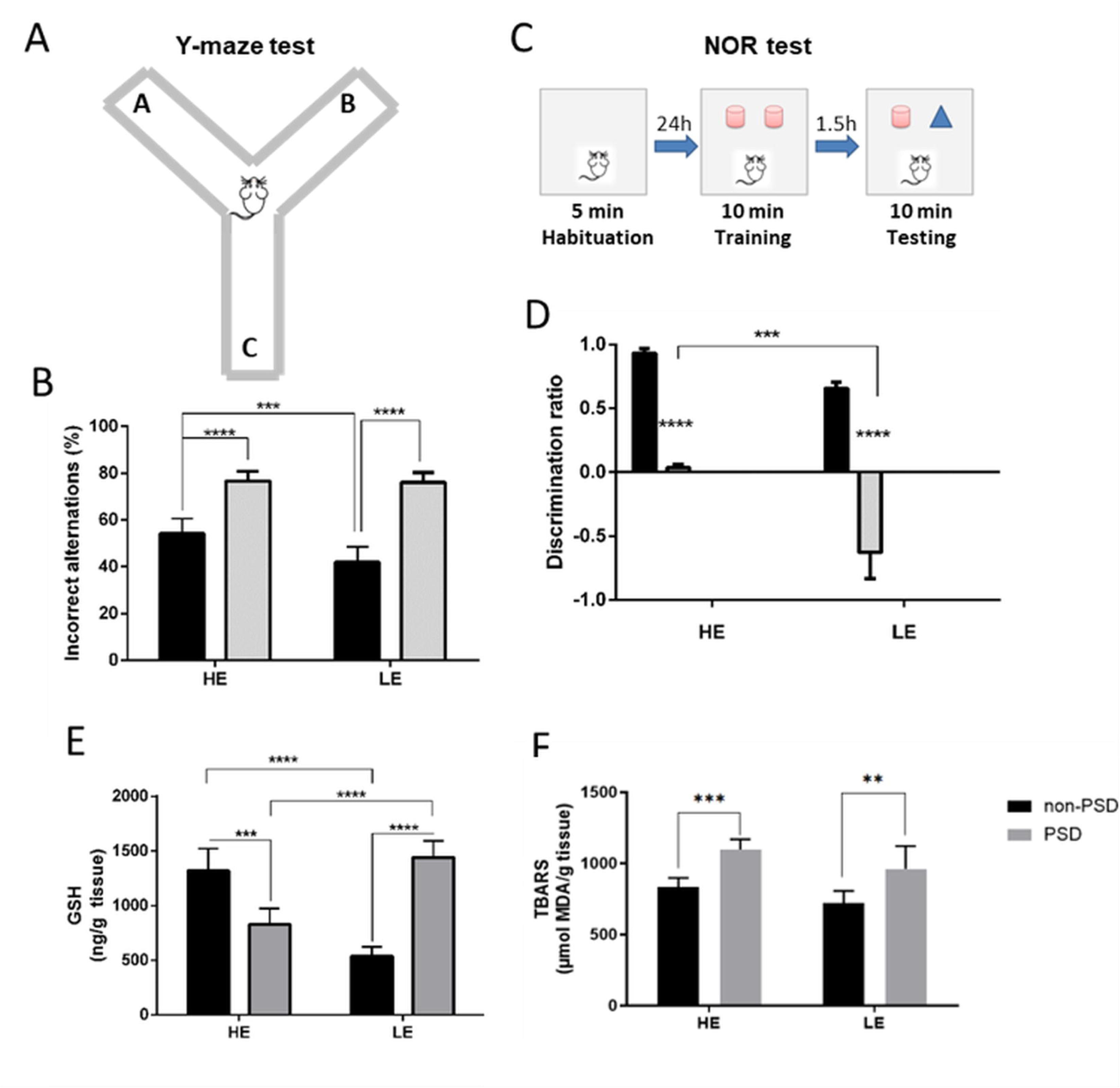
**Cognitive performance and oxidative stress in High-exploratory (HE) and Low-exploratory (LE) rats: Effects of paradoxical sleep deprivation (PSD) on working memory and redox status** (A) Y-maze testing apparatus. (B) Percentage of incorrect alternations in the Y-maze test, showing increased errors in the HE-PSD and LE-PSD group compared to their respective control groups. (C) Novel Object Recognition (NOR) testing apparatus. (D) The discrimination ratio in the NOR test shows enhanced cognitive performance in the HE groups compared to the LE group. However, both HE and LE rats in the PSD groups exhibited a significant decline in cognitive performance when compared to their non-PSD counterparts. (E) Tissue levels of GSH (Glutathione) in the hippocampus showed a significant reduction in the HE-PSD group compared to its control, while a significant increase was observed in the LE-PSD group compared to its control group. (F) TBARS (Thiobarbituric Acid Reactive Substances) levels in hippocampal tissue, reflecting oxidative stress, showed significant increases in both the HE-PSD and LE-PSD groups compared to their respective controls. *p < 0.05, **p < 0.01, ***p < 0.001, ****p < 0.0001. Data were analyzed using appropriate statistical tests. The results indicate that PSD induces significant changes in cognitive function and oxidative stress, with variations observed based on exposure levels and PSD status.

Y-maze results showed increased incorrect alternations in HE+PSD and LE+PSD rats compared to non-PSD controls (Figure 5B, P < 0.0001). Additionally, HE-non-PSD rats exhibited more incorrect alternations than LE-non-PSD rats (P = 0.001).

In the NOR test, PSD reduced the discrimination ratio [F(1,19) = 4.190, P = 0.0548], with HE+PSD and LE+PSD rats showing significant impairments relative to non-PSD controls (P < 0.0001). LE+PSD rats displayed a more pronounced reduction than HE+PSD rats (Figure 5D, P = 0.0001).

Hippocampal GSH levels exhibited a significant interaction between “Exploratory phenotype” and “PSD exposure” [F(1,19) = 130.4, P < 0.0001]. HE-non-PSD rats showed higher GSH levels than LE-non-PSD (P = 0.0001). PSD reduced GSH in HE+PSD (P = 0.0003) but increased it in LE+PSD (P < 0.0001) relative to controls. Notably, GSH levels in LE+PSD rats were significantly higher than in HE+PSD (P < 0.0001) (Figure 5E). Lipid peroxidation (Figure 5F) was elevated in HE+PSD (P = 0.0005) and LE+PSD (P = 0.0017) compared to controls.

### 3.5 Hippocampal Inflammatory and Neuroplasticity Alterations Induced by PSD

For hippocampal nitrite and iNOS levels, two-way ANOVA revealed a significant interaction between “Exploratory phenotype” and “PSD exposure” [F(1,22) = 4.733, P = 0.0406; F(1,11) = 6.616, P = 0.0260]. Nitrite levels increased in PSD-exposed HE and LE rats compared to controls (HE+PSD vs. HE-non-PSD, P < 0.0001; LE+PSD vs. LE-non-PSD, P = 0.0003) (Figure 6A). Given the observed nitrite differences, iNOS expressions were analyzed. PSD significantly increased iNOS mRNA levels in HE+PSD rats compared to HE-non-PSD (P = 0.0022) (Figure 6B).

**Figure 6.**
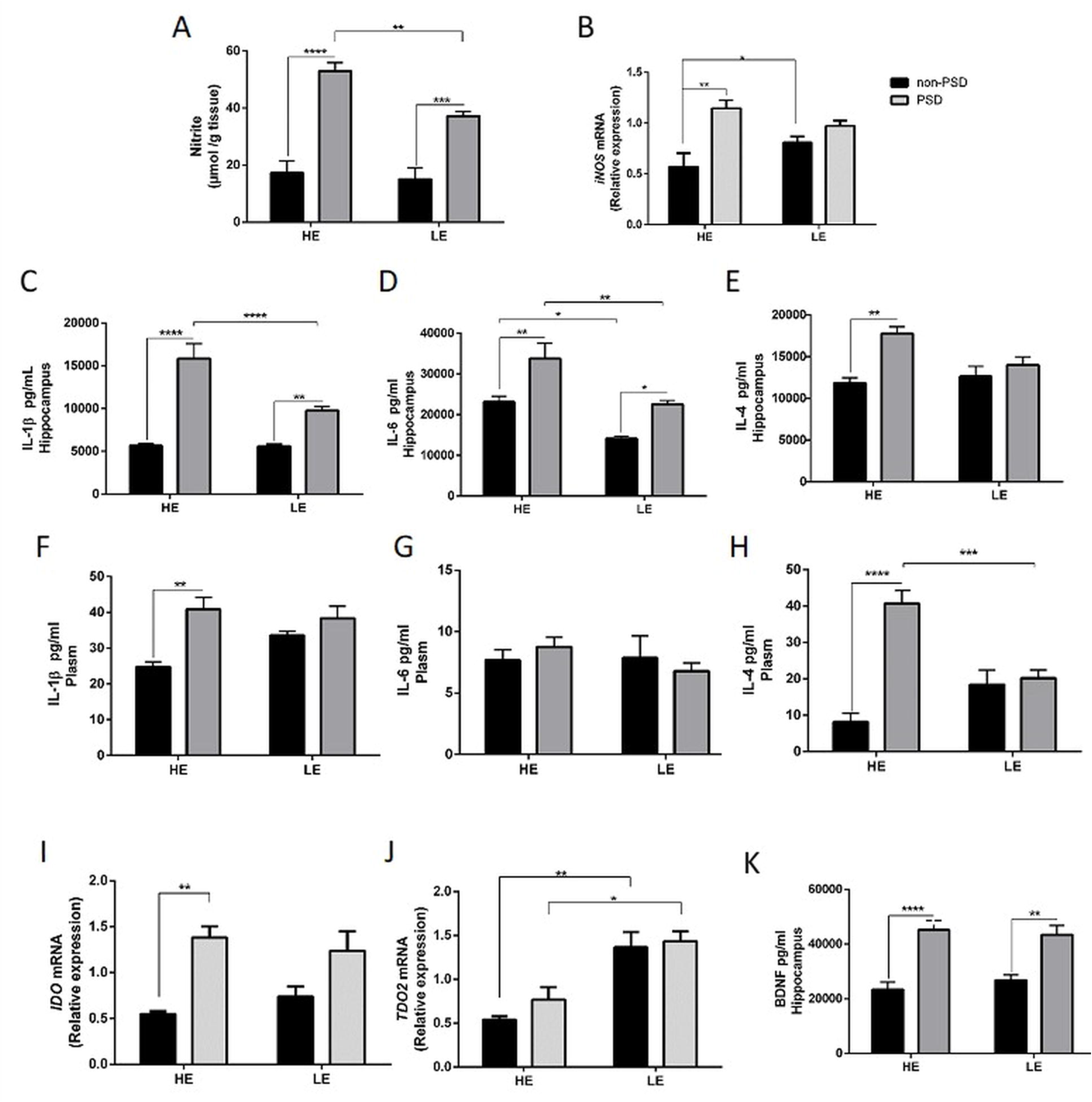
**Inflammatory and molecular markers in High-exploratory (HE) and Low-exploratory (LE) rats: effects of paradoxical sleep deprivation (PSD) on cytokines, nitrite levels, and gene expression.** Levels of hippocampal nitrite (A), IL-1β (C), IL-6 (D), IL-4 (E) cytokines and BDNF (K); plasma cytokines IL-1β (F), IL-6 (G), IL-4 (H) by ELISA; and relative *m*RNA levels of *iNOS* (B) *IDO1* (I), *TDO2* (J) by qPCR assay in the hippocampus of the HE- and LE-rats exposed to chronic PSD for 10 days. Each bar represents the mean ± S.E.M of 8–10 animals/group. Data were analyzed by one-way ANOVA, with Tukey as a post hoc test. *P < 0.05, **P < 0.01, ***P < 0.001, ****P < 0.0001. IL-1β: Interleukin-1 beta, IL-6: Interleukin-6, IL-4: Interleukin-4, iNOS: Inducible Nitric Oxide Synthase, IDO: Indoleamine 2,3-Dioxygenase, TDO: Tryptophan 2,3-Dioxygenase.

A significant interaction between “Exploratory phenotype” and “PSD exposure” was observed in hippocampal IL-1β [F(1,30) = 15.02, P = 0.0005], IL-6 [F(1,29) = 0.3274, P = 0.5716], and IL-4 levels in both hippocampus and plasma [F(1,27) = 4.860, P = 0.0362; F(1,22) = 25.69, P = 0.0001]. Post-hoc analysis showed increased hippocampal IL-1β in HE+PSD and LE+PSD compared to controls (P < 0.0001 and P = 0.0042, respectively) (Figure 6C). Plasma IL-1β was elevated in HE+PSD vs. HE-non-PSD (P = 0.0078) (Figure 6F).

IL-4 levels were significantly higher in the hippocampus of HE+PSD compared to HE-non-PSD (P = 0.0031) (Figure 6E). Plasma IL-4 was also increased in HE+PSD vs. HE-non-PSD (Figure 6H). IL-6 levels were elevated in HE+PSD and LE+PSD rats relative to controls (HE: P = 0.0024; LE: P = 0.0323) (Figure 6D).

For *Ido*1 gene expression, a two-way interaction was observed [F(3,12) = 1.579, P = 0.2329], with increased *Ido*1 mRNA in HE+PSD vs. HE-non-PSD (P = 0.0040) (Figure 6I). *Tdo*2 expression was higher in LE+PSD vs. HE+PSD (P = 0.0267) and in LE-non-PSD vs. HE-non-PSD (P = 0.0081) (Figure 6J).

BDNF levels were elevated in HE+PSD vs. HE-non-PSD (P < 0.0001) and in LE+PSD vs. LE-non-PSD (P = 0.0018) (Figure 6K).

### 3.6 PSD Modulates Circadian Clock Gene Expression in HE and LE Rats

The expression of key circadian clock genes was significantly altered by PSD exposure, with distinct effects observed in HE and LE rats. A Student’s t-test revealed significant differences in the expression of *Bmal1*, *Clock*, *Per1*, *Per2*, *Per3*, *Cry1*, and *Cry2* (Figure 7A-G).

**Figure 7.**
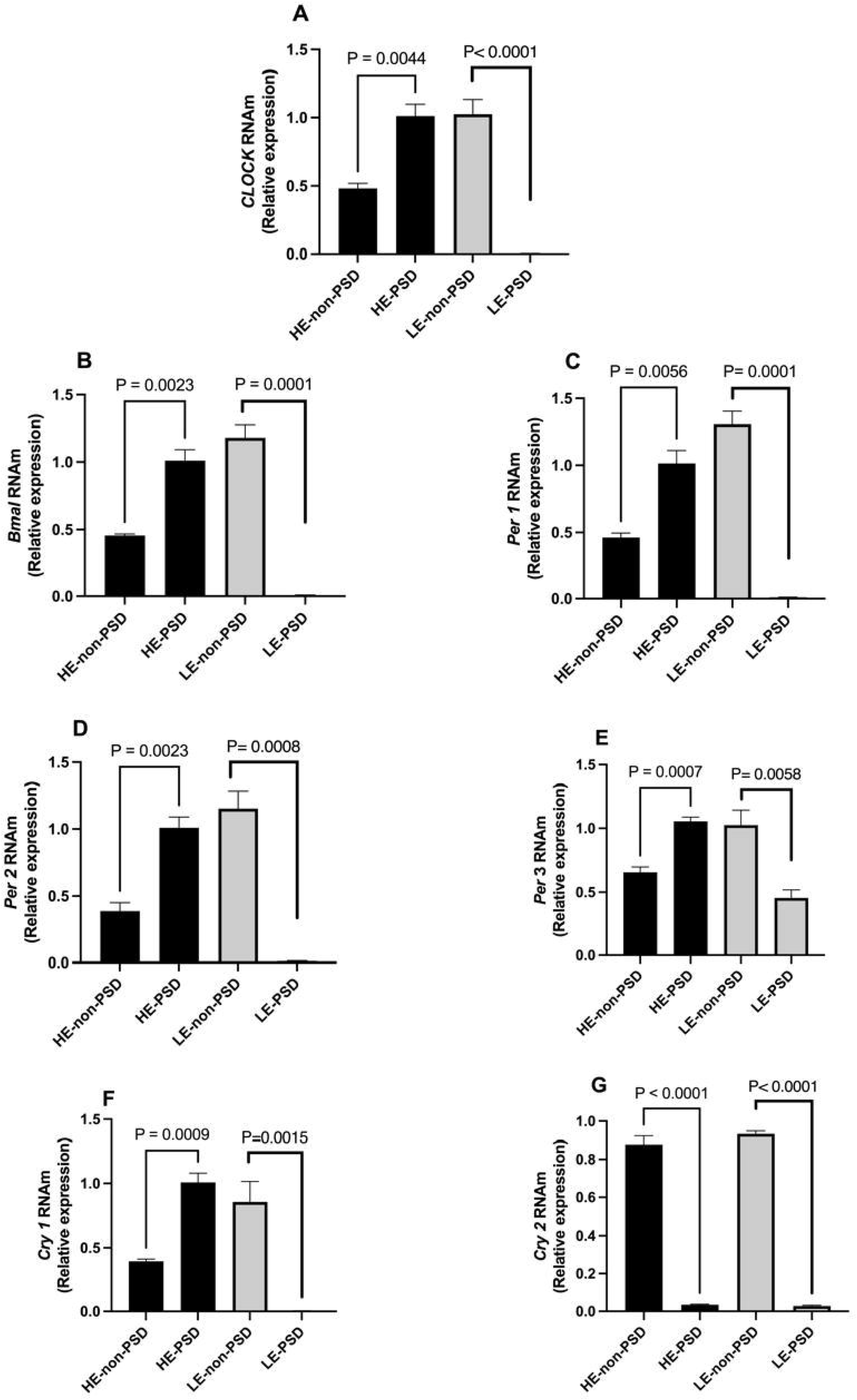
**PSD alters the expression of circadian clock genes in High-exploratory (HE) and Low-exploratory (LE) rats.** *m*RNA expression levels of *Bmal1* (A), *Clock* (B), *Per1* (C), *Per2* (D), *Per3* (E), *Cry1* (F), and *Cry2* (G) were analyzed in the hippocampus of HE and LE rats exposed to PSD or control conditions. Data are presented as mean ± SEM. Statistical comparisons were performed using Student’s t-test. P < 0.05 was considered statistically significant. Clock: Circadian Locomotor Output Cycles Kaput, Bmal1: Brain and Muscle ARNT-Like Protein 1, Per1: Period Circadian Regulator, Per2: Period Circadian Regulator 2, Per3: Period Circadian Regulator 3, Cry1: Cryptochrome Circadian Regulator 1 and Cry2: Cryptochrome Circadian Regulator 2.

HE-non-PSD rats exhibited higher basal expression levels of *Clock*, *Bmal1*, *Per1*, *Per2*, *Per3*, and *Cry1* compared to LE-non-PSD rats.

Following PSD exposure, HE+PSD rats showed an upregulation of *Clock* (P = 0.0044), *Bmal1* (P = 0.0023), *Per1* (P = 0.0056), *Per2* (P = 0.0023), *Per3* (P = 0.0007), and *Cry1* (P = 0.0009) relative to HE-non-PSD rats. Conversely, in LE+PSD rats, PSD significantly reduced the expression of *Clock* (P < 0.0001), *Bmal1* (P < 0.0001), *Per1* (P = 0.0001), *Per2* (P = 0.0008), *Per3* (P = 0.0058), and *Cry1* (P = 0.0015) compared to LE-non-PSD.

Additionally, *Cry2* expression was significantly decreased in both HE-PSD and LE-PSD groups (P < 0.0001 for both) compared to their respective controls. These findings suggest a differential regulation of circadian rhythm genes in HE and LE rats in response to PSD exposure, with HE rats exhibiting an upregulation and LE rats showing a downregulation in gene expression.

### 3.7 Chronic PSD Differentially Modulates D1 and D2 Expression in HE and LE Rats

Chronic PSD exposure increased hippocampal D1 expression in HE+PSD compared to HE-non-PSD rats (P < 0.001, Figure 8A). Significant differences were also observed between HE+PSD and LE+PSD rats (P < 0.001, Figure 8A). In contrast, D2 expression was only elevated in LE+PSD compared to HE+PSD rats (P < 0.05, Figure 8B).

**Figure 8.**
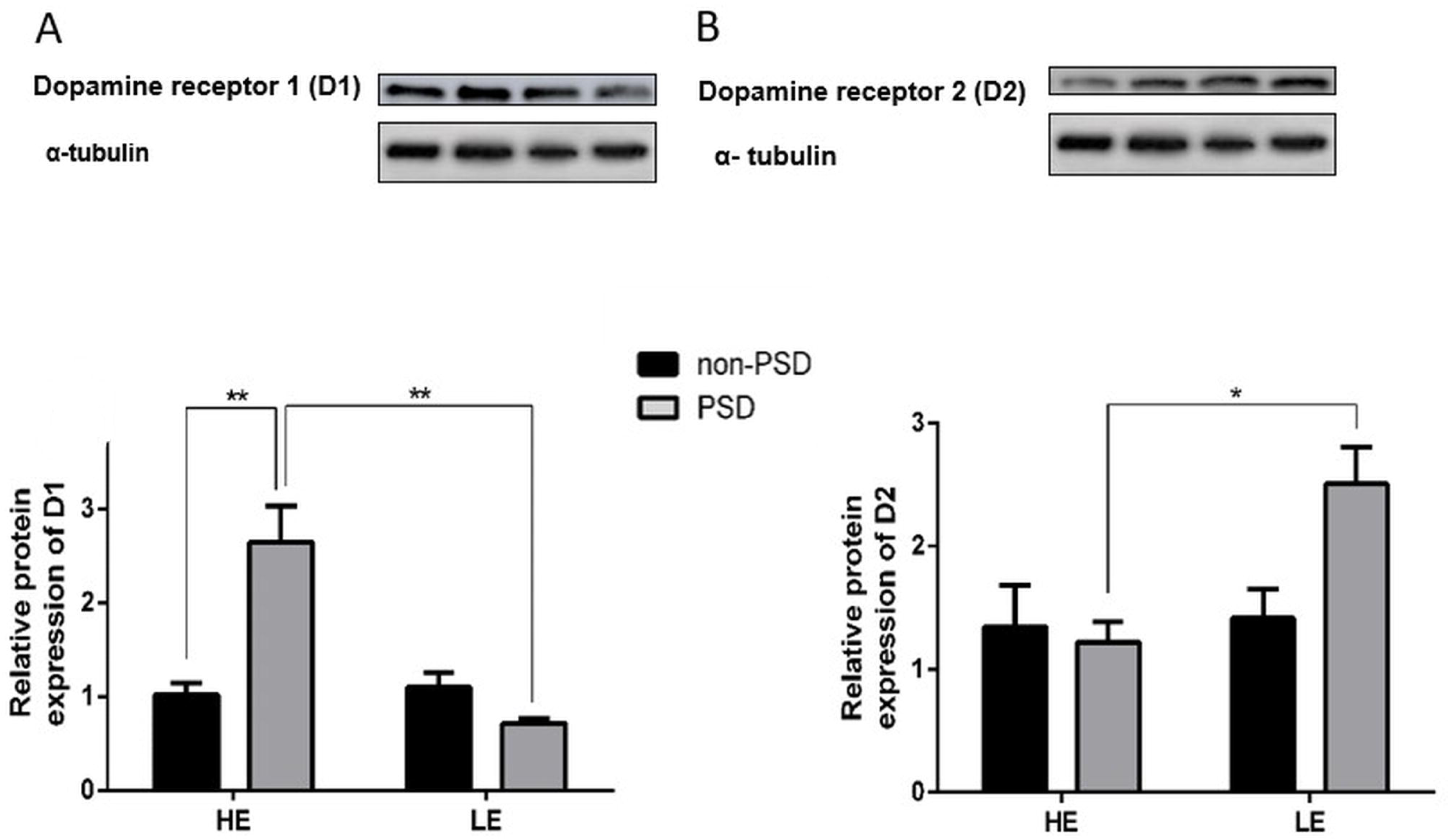
**Dopamine receptor expression in High-exploratory (HE) and Low-exploratory (LE) rats: Effects of paradoxical sleep deprivation (PSD) on D1 and D2 protein levels** Hippocampal relative *m*RNA levels of D1(A) and D2 (B) by qPCR assay in rats HE and LE exposed to chronic PSD (10 days). All fold changes were calculated by the ΔΔCt method. Each bar represents the mean ± S.E.M of 8–10 animals/group. Data were analyzed by one-way ANOVA, with Tukey as a post hoc test. *P < 0.05, **P < 0.01. D1: Dopamine receptor 1, D2: Dopamine receptor 2, qPCR: Quantitative polymerase chain reaction.

## 4. DISCUSSION

Our study demonstrated that PSD differentially affects behavior, neuroinflammation, oxidative stress, and circadian regulation in HE and LE rats, highlighting phenotype-dependent stress susceptibility and resilience mechanisms. HE rats exhibited higher uric acid levels, increased risk-taking and exploratory behavior, and greater circadian gene upregulation (*Bmal1, Clock, Per1-3, Cry1*) following PSD. In contrast, LE rats showed greater vulnerability to depressive-like behaviors, including reduced sucrose preference, increased immobility, and downregulation of circadian genes. Oxidative stress responses also varied, with HE rats showing higher basal GSH, which was depleted by PSD, whereas LE rats exhibited a PSD-induced GSH increase, suggesting differential redox resilience.

Additionally, both phenotypes exhibited heightened neuroinflammation, with increased nitrite, iNOS, IL-1β, and IL-6, though IL-4 was selectively elevated in HE rats, indicating a potential anti-inflammatory adaptation. Lastly, D1 and D2 were altered by PSD, further linking dopamine-circadian interactions to mood regulation. Together, these findings emphasize how individual phenotypes shape behavioral and molecular responses to chronic stress, providing insights into mechanisms of stress vulnerability and resilience.

Our findings reveal that uric acid levels are associated with impulsivity-related behaviors, with HE rats exhibiting higher uric acid concentrations and greater exploratory behavior, whereas LE rats showed increased avoidance behavior negatively correlated with uric acid. These results align with previous studies linking uric acid to impulsivity in clinical and nonclinical populations (Albert et al., 2015; Manowitz et al., 1993; Sutin et al., 2013), suggesting a potential role of purinergic metabolism in modulating risk-taking behavior.

Additionally, PSD differentially affected exploratory behavior, anxiety, depression, and anhedonia in HE and LE rats. While HE rats exhibited increased risk-taking and exploratory behavior following PSD, LE rats displayed greater vulnerability to depressive-like behaviors, indicating that LE rats might be more prone to mood disturbances under sleep deprivation (Katz, 1982; Mällo et al., 2007; Stedenfeld et al., 2011a, 2011b; Veenema et al., 2003). These findings suggest that individual differences in exploratory phenotypes influence susceptibility to PSD-induced mood alterations.

Changes in *Tph2 m*RNA levels further support the notion of altered serotonergic activity contributing to these behavioral outcomes (Lieb et al., 2019). Increased expression of the *Tdo2* gene and reduced expression of the *Tph2* gene in the brain suggest significant alterations in tryptophan metabolism, which can have implications for serotonin synthesis and overall neurochemical balance (Chen and Miller, 2012; Hattori et al., 2018; Kanai et al., 2009; Markett et al., 2017). The combination of increased *Tdo2* and reduced *Tph2* expression suggests a scenario where tryptophan is preferentially metabolized via the kynurenine pathway at the expense of serotonin synthesis (Höglund et al., 2019; Miura et al., 2008) .This imbalance in tryptophan metabolism could be associated with changes in mood, behavior, and cognitive function, potentially influenced by stress, inflammation, or other physiological stressors (Allison and Ditor, 2014; Karu et al., 2016; O’Connor et al., 2021; Parrott et al., 2016).

Our study also demonstrates that PSD impairs working and declarative memory, particularly in LE rats, while also altering oxidative stress responses. The behavioral heterogeneity of Wistar rats influences data dispersion in behavioral evaluations (de Oliveira-Júnior et al., 2024; Palm et al., 2011; Redei et al., 2023; Theilmann et al., 2016). Therefore, we suggest associating the PSD model with the exploratory standard (temperamental base), as temperament can modify individual vulnerability to mood disorders (Middeldorp et al., 2011; Simonetti et al., 2023). Temperaments may carry a genetic influence (Cloninger et al., 1993b; Smoller and Finn, 2003). Selecting animals based on exploratory standards can facilitate modeling natural neurobiological substrates and potential, representing temperament differences and possibly carrying a genetic predisposition to different mood disorders.

HE rats exhibited higher basal GSH levels, which were significantly reduced by PSD, whereas LE rats showed an increase in GSH after PSD exposure. This suggests differential oxidative stress resilience mechanisms, with HE rats being more resistant to oxidative insults but more sensitive to PSD-induced GSH depletion.

Furthermore, PSD-induced hippocampal neuroinflammatory alterations were observed in both HE and LE rats, with increased nitrite levels, iNOS expression, and pro-inflammatory cytokines (IL-1β and IL-6). Interestingly, IL-4 levels were selectively elevated in HE rats, suggesting a compensatory anti-inflammatory response. Elevated pro-inflammatory cytokines (IL-1β, IL-6) and BDNF levels suggest an inflammatory response to PSD, contributing to behavioral and cognitive alterations (Khadrawy et al., 2011; Rosa Neto et al., 2010).

We also found that circadian clock genes expression was differentially regulated by PSD, with HE rats exhibiting upregulation and LE rats showing downregulation of key circadian genes (*Bmal1, Clock, Per1-3, and Cry1*). These changes were accompanied by PSD-induced alterations in hippocampal D1 and D2 expression, further supporting the role of dopaminergic signaling in modulating circadian disruptions and mood-related behaviors. PSD is capable of inducing essential changes in the dopaminergic system, with hyperdopaminergic states linked to mania alterations (Frey et al., 2006; Young et al., 2010). Previous studies have reported that *Clock* gene manipulation, specifically within the mesolimbic dopamine system, produced animals with a behavioral phenotype similar to HE+PSD (Roybal et al., 2007b). Our results indicate that *Clock* gene expression and D1 protein levels increased in the hippocampus of HE+PSD rats. This aligns with studies showing transcriptional changes in the *Clock* and *Per1* genes in the mesolimbic system after treatment with cocaine or methamphetamine (Michihiko et al., 2002; Nikaido et al., 2001; Uz et al., 2005; Yuferov et al., 2003).

In contrast, LE+PSD rats showed a behavior profile similar to depression and anhedonia. This genetic LE+PSD profile is similar to mice exposed to chronic moderate stress, which produced chronic clock gene changes, such as decreased heterodimer *Clock/Bmal1* and reduced expression of *Cry2, Per1, and Per2* (Calabrese et al., 2016). These gene changes could be related to a link between stress exposure and clock gene dysfunction, represented by changes in HPA axis function (Buijs and Kalsbeek, 2001). As mentioned above, we found that LE+PSD rats reduced tryptophan hydroxylase 2 (*Tph2*) gene expression, suggesting reduced serotonin synthesis, which plays a role in depression pathophysiology (Cowen and Browning, 2015).

Our results, along with previous studies, indicate differential activation of D1 in HE rats and D2 in LE rats (Easton et al., 2003; Freyberg and McCarthy, 2017; Hood et al., 2010; Imbesi et al., 2008; Zhang et al., 2021); however, the mechanisms by which dopamine and D1- or D2-activation positively or negatively regulate *Clock* gene expression in the hippocampus remain to be resolved. Future experiments are needed to elucidate if PSD-induced changes in *Clock* gene expression are responsible for differential behavior regulation in these animals.

The main limitation is species differences, as rats have lower uric acid levels than primates, limiting generalizability. Additionally, while associations between clock genes and dopaminergic alterations were observed, causal mechanisms remain unclear. Despite this, the study provides key insights into temperament and neuropsychiatric disorders, supporting future cross-species research.

Overall, our results highlight the complex interplay between exploratory phenotype, oxidative stress, neuroinflammation, and circadian gene regulation in response to PSD, providing insights into individual differences in stress susceptibility and resilience mechanisms.

## Supporting information

Supplemental methods and Results

## Acknowledgments

We thank the Brazilian National Council for Scientific and Technological Development (CNPq) for financial support through a doctoral fellowship. We also acknowledge CAPES and FUNCAP for their support with scholarships.

## Declaration of generative AI and AI-assisted technologies in the writing process

During the preparation of this work, the authors used CHAT GPT 4o to revise the manuscript grammar to improve readability. After using this tool/service, the authors reviewed and edited the content as needed and take full responsibility for the content of the publication.

