## Supplemental methods and Results for "The Sleep-Mood Connection: Temperament Modulates Neuroinflammation, Clock Genes, and Dopaminergic Receptors Expression in Rats"

### Supplementary Methods and Materials

#### Animals

The experiments were performed in Male *Wistar* periadolescent (weighting 50–60 g) obtained from the Animal House of Universidade Federal do Ceará. The animals were housed at the maximum of 5 per cage in standard polycarbonate cages (42 × 20.5 × 20 cm) and standard environmental conditions (22 ± 1 °C; humidity 60 ± 5%; 12-h light/dark cycle with lights on at 7:00 a.m.) with access to food (Laboratory RodentDiet - LabDiet®) and water ad libitum. All behavioral procedures were conducted between 8:00 and 14:00 h by raters blinded to the experimental groups. The methods followed the guidelines outlined in Brazilian regulations and at the NIH Guide for the Care and Use of Laboratory Animals and were approved by the local Ethics Committee for Animal Research (CEPA) at Universidade Federal do Ceará under protocol number 125/14. All efforts were made to minimize animal suffering and reduce the number of animals used in the study.

#### Behavioral analyses of HE- and LE- rats

#### Screening for locomotor response to novelty

A psychogenic selection was employed to study the bases of temperament in rodents, dividing them into extremes of exploratory activity (1,2). To achieve this, peri-adolescent animals on the 30th postnatal day (PND) were selected based on their performance during a 10-minute session in the open field test. The rats were then divided into two groups to represent extremes of temperament: HE rats and LE rats, categorized according to their 25$^{th}$ percentile and 75$^{th}$ percentile vertical exploratory activity levels (3) (Figure 1). These animals were tested again in this same apparatus in adult life and maintained the same exploratory profile. All testing was performed between ZT2 (8 a.m.) and ZT5 (11 a.m.)

After screening 80 rats in the task involving reaction to novelty (percentage of rearing in the open field) in a new environment, the 20 top and 20 bottom rats regarding rearing behavior in the open field were selected and defined, respectively, as HE and LE, corresponding, therefore, to extremes of temperament. Animals in the 75th percentile or higher were classified as HE, as they exhibited a greater percentage of vertical exploratory behaviors compared to those in the 25th percentile or lower, who were classified as LE [HE 80.60 ± 9.992 %, N=20); (LE 21.90 ± 7.549%, N= 20); t = 21.29, df=39, p <0.0001]. The upper and lower cut-off values for classification were greater than 67.25% for the HE groups and less than 32% for the LE group, corresponding to the 75th and 25th percentiles, respectively (Supplementary Table 1). The remaining 39 animals with a rearing percentage between 32% and 67.25% in the open field were considered median and did not enter the study. The same task was performed one month later to confirm that these behavioral traits were reproducible (PND60). Indeed, the two selected groups remained very distinct (t = 8.254; P < 0.0001) in the second trial.

**Experimental Protocol**

After the distinction of HE and LE rats on PND30, the animals from PND40-49 were randomly selected to be exposed or not to PSD. For this end, animals were submitted to a ten-day protocol with PSD exposure on alternative days (see Figure 1). For PSD sessions, the rats were placed in 5 per cage (41 cm × 34 cm × 16.5 cm), each cage containing 12 platforms (3 cm diameter). In the same box, we put a volume of water 1 inch deep, obligating the animals to stay on the platforms. The animals were allowed to move freely between platforms. During the paradoxical sleep phase, characterized by muscle atonia, they were awakened by falling into the water. Food and water were available ad libitum(4). This study employed 24 hours of paradoxical sleep deprivation (PSD) as it has been shown in previous studies to increase locomotor activity, a behavior indicative of mania-like symptoms(5). Control group rats were subjected to the same conditions, except no water was present at the bottom of the box.

Based on this protocol, we established four experimental groups (n = 8/group): Group 1—HE-control (non-stressed), Group 2—HE+PSD, Group 3—LE-control (non-stressed), and Group 4—LE+PSD. The rats were subjected to a series of tests described below: the aversive exposure to cat odor test to evaluate risk-taking behavior, the open-field test for assessing locomotion, the elevated plus maze for anxiety assessment, the forced swimming test for depression-like behavior, the sucrose preference test for evaluating anhedonia, the Y-maze test for assessing working memory, and the novel object recognition test for evaluating declarative memory. Following euthanasia, the rats' blood or brain tissue were examined for uric acid levels, reduced glutathione (GSH) levels, lipid peroxidation, nitrite determination, and were subjected to immunoassays for IL-1β, IL-4, and IL-6, as well as gene expression analysis of *CLOCK, Bmal1*, *Cry1, Cry2, Per1, Per2, and Per3*, and protein expression analysis of D1- and D2-like dopaminergic receptors.

### Behavioral parameters analyzed

#### Aversive exposure to cat odor test

On the day of exposure, rats were individually placed in a box for 10 minutes, alongside either a cat odor or a neutral odor block. One hour before the experimental session, cat odor was prepared by vigorously rubbing a damp cloth (18 × 22 cm) against the skin of an adult female domestic cat for five minutes. The cat-odor cloth was stored in a sealed plastic bag and was used for a maximum of four exposures. Portions of the original cloth that were not rubbed on the cat were used to create a neutral odor. All odor exposures were conducted in a separate, small, dimly lit room. To prevent contamination, neutral odor exposures always preceded cat odor exposures. Before the first cat odor exposure, the cloth was placed in the test room for 10 minutes to acclimate. Each rat was transported to the exposure room in its home cage, positioned next to the odor cloth wedged between the cage tops. The cloth was placed at the end opposite the food and water containers (Figure 2D). The test duration was 10 minutes, during which rats were videotaped for subsequent analysis. The frequency and duration (in seconds) of cloth contacts were defined as direct contact or sniffing within 5 cm of the cloth. The frequency and duration of sheltering were recorded when a rat was underneath the food and water compartments (6).

**Open-field test (OFT)**

The open-field area was made of acrylic (with walls and a black floor, 50 X 50 cm) divided into five squares of equal areas, an exploratory activity of the animal was recorded for 10 minutes and the trajectory traversed by each animal in the arena. We investigated the following parameters: time spent in seconds without central quadrant; the relationship between a locomotor activity in the central quadrant and the quadrants of the arena periphery (C / P ratio); the immobility time in seconds (s) and the number of transitions between the quadrants of the arena. Besides, the number of rearing (number of clothing or animal raised on the hind legs or vertical locomotive activity) was counted through observation, regardless of whether or not used to support animals on the walls (Figure 3A). The test was performed in a room with attenuated sound, in the low light intensity condition, recorded and analyzed, using the software SMART video tracking version 3.0.03 of Panlab Harvard Apparatus®.

**Elevated plus maze (EPM) test**

This test is traditionally used to assess anxiety-like behavior (7). Twenty-four hours following the final open field exposure, animals were evaluated using the elevated plus-maze. Constructed from black-painted Plexiglas, the maze features two opposing open arms (50 x 10 cm) and two opposing closed arms, each enclosed by 40 cm high walls. The maze was elevated 50 cm above the floor. Behavioral testing was conducted under red light conditions with a 68 dB wide spectrum masking noise present. Each rat was placed in the central square facing an open arm. The number of entries into each arm (indicated by all four paws entering the arm) and the time spent in each arm were recorded for 5 minutes (Figure 4A). For analysis, open-arm activity was quantified by calculating the percentage of entries into the open arms relative to the total number of entries into all arms (open entries/total entries × 100), as well as the percentage of time spent in the open arms relative to the total time spent in the maze (open arms/total time × 100), and the percentage of time spent in the closed arms relative to the total time spent in the maze (closed arms/total time × 100).

**Forced swimming test (FST)**

The FST was first described by Porsolt et al. (8,9). This model is based on the evaluation of immobility as a measure of behavioral despair, which rodents adapt to after being placed in a condition from which they cannot escape. Rats were individually forced to swim into a vertical glass cylinder (22.5 cm diameter and height 60 cm) containing about 35 cm water at 25 ◦C. On the first day of the experiment, the procedure lasted 10 min and the re-exposition 24 h later lasted 5 min, and the total time spent immobile was recorded (10). The rat was judged to be immobile when it remained floating in the water with all limbs motionless and to make only very minimal movements necessary to keep its head above the water (Figure 4E). The water was changed after every trial. At the end of each session, the rats were dried with laboratory tissues. An increase in the duration of immobility is indicative of depressive-like behavior.

**Sucrose preference test**

Sucrose preference tests were used to define anhedonia operationally. Specifically, anhedonia was described as a reduction in sucrose intake and sucrose preference relative to the intake and preference of the control group(11). A sucrose preference test consisted of first removing the food and water from each rat cage (both PSD and control groups) for 20 h. Drinking water and 1% w/v sucrose were then placed in the cages in preweighed glass bottles, and animals were allowed to consume the fluids freely for a period of 1 h (Figure 4G). Sucrose preference was measured by calculating the proportion of sucrose consumption out of the total consumption of liquid. The test was carried out on two consecutive days.

**Y-maze test**

This test was used to assess spatial working memory by spontaneous alternation performance, which allows the evaluation of cognitive searching behavior(12). A Y-maze apparatus made of black acrylic consisted of three arms with 425 mm (length), 145 mm (width), and 225 mm (height) mounted symmetrically (120° between arms) to an equilateral triangular center compartment. Each rat was placed at the end of one arm and allowed to freely move through three arms of the maze for 8 min. The series of arm entries was recorded visually (Figure 5A). The number of maximum alternations was the total number of arms entered minus 2, and the percent alternation was calculated as a total of alternations / (total arm entries − 2) X 100(13). An alternation was considered correct if the animal visited a new arm and did not return to a previously visited arm (example of correct alternation: 1, 2, 3 arms; an example of incorrect alternation: 1, 2, 1 arm).

**Novel object recognition test (NOR)**

This test is used to evaluate the preference for a new object-related in rodents to declarative memory (14). On the first day, before any procedure rats were habituated, the animals were put in the center of the open field and left for 10 min with no object. After 24 h (training session), animals were exposed to two identical copies of objects A1 and A2 (double Lego® toys) for 10 minutes to explore the new environment, and the exploration time was registered. Rats were then removed from the open field for 1, 5 hr, during which the initial objects were replaced by an identical copy of object A1 and a novel object B1 in color and size, but of different shapes. Rats were then reinserted into the open field and left to explore for 10 min (15,16) (Figure 5C). The time spent in the direct exploration of the objects was recorded for further analysis. Objects and the arena were cleaned with 50 % ethanol and water between animals. The identity of the first object and its position in the field were counterbalanced. The recognition index for the novel object was calculated as the ratio of the time spent with the novel object divided by the time spent with both objects combined (17).

#### 2.6. Determination of oxidative and inflammatory parameters

##### 2.6.1 Uric Acid levels

Uric acid levels were routinely monitored in serum samples from the rats using a commercial kit (Randox, San Diego, California).

##### 2.6.2. Assays for Oxidative Stress Determinations

The GSH content of the brain homogenate was measured at 412 nm using the method of Ellman's reagent (DTNB) reaction with free thiol groups (18). Briefly, the supernatants were mixed with 0.4 M Tris–HCl buffer, pH 8.9, and 0.01 M DTNB. Reduced glutathione levels were determined by the absorbance at 412 nm and were expressed as ng of GSH/g wet tissue.

Lipid peroxide levels in brain homogenate were measured with the thiobarbituric-acid reactive substances (TBARS) as an index of reactive oxygen species (ROS) production by the method of Placer et al (19). The quantification of TBARS was determined by comparing the absorption to the standard curve of malondialdehyde (MDA) equivalents generated by acid-catalyzed hydrolysis 1,1,3,3 tetra methoxy propane. Lipid peroxidation was estimated by the absorbance at 532 nm and expressed as μmol of MDA/mg of tissue.

Nitrite determination to assess alterations in nitric oxide (NO) production was determined in the rat brain homogenates. The production of NO was determined based on the Griess reaction (20,21). The absorbance was measured at 550 nm via a microplate reader. The standard curve was prepared with several concentrations of NaNO2 (ranging from 0.75 to 100 μM) and was expressed as μmol/g of protein.

##### 2.6.3 Immunoassay for IL-1β, IL-4, IL-6

The brain areas were homogenized in 8 volumes of PBS buffer with protease (EMD Biosciences) inhibitor and centrifuged (10,000 rpm, 5 min). The concentration of the cytokines in 50 ml samples was determined by the immunoenzymatic assay ELISA (R&D Systems, Minneapolis, MN, USA), according to the manufacturer's protocol, and expressed in pg/g tissue.

### 2.7 RNA isolation for gene expression analyses

Total RNA was isolated using an RNA isolation protocol (PROMEGA). RNA was quantified by NanoDrop (Thermo Fisher Scientific), and RNA quality was determined by examining the 260/280 ratio > 1.8. A total of 1 µg RNA was then reverse-transcribed using a High-capacity cDNA reverse transcription Kit (Applied Biosystems) according to the manufacturer protocol. *m*RNA expression was analyzed by quantitative PCR (qPCR) according to the manufacturer's instructions using primers (Applied Biosystems, USA). The sequences of primers ID numbers are listed in Supplementary Table 2**.** Target gene expression is calculated relative to a stably expressed reference gene (GAPDH gene). A total of 50 ng cDNA was added to the gene expression assay (1 µL *Clock, Bmal1, Cry1, Cry2, Per1, Per2 and Per 3* or G*apdh*), Power SYBR Green PCR master mix (10 µL), and RNA-free water (5 µL) to a final volume of 20 µL. After the reaction components were mixed by inverting the tube several times, the tube was briefly centrifuged. Then, 20 µL of the PCR reaction mix was transferred to each well. The plate was loaded into the instrument, sealed, and centrifuged. The thermocycler parameters were 50°C for 2 min and 95°C for 10 min, followed by 40 cycles of 95°C for 15 s and 60°C for 60 s. All fold changes were calculated by the $\Delta\Delta C_t$ method.

### 2.8 Western Blotting

Hippocampus were homogenized in RIPA lysis buffer (25 mM Tris-HCL, pH 7.6; 150 mM NaCl; 5 mM EDTA; 1% NP40; 1% Triton X-100; 1% sodium deoxilato; 0,1% SDS) and protease inhibitor (1µL inhibitor: 100 µL RIPA). To protein extraction, Hippocampi samples were centrifuged (17 min, 4ºC, 13000 rpm), and the supernatant was collected. Protein concentrations were determined through the method of Bradford, according to the manufacturer of the protocol. SDS-polyacrylamide gel electrophoresis (10%) was performed using 20 µg of protein (previously prepared with Laemmli sample buffer- and heated at 95ºC for 5 minutes). After, the protein was transferred to PVDF membrane, blocked with BSA 5% for 1 hour, incubated overnight with primary antibody (rabbit anti-D1, rabbit anti-D2, or mouse anti-tubulin) and secondary antibody (Goat anti-rabbit or Goat anti-Mouse IgG). According to the manufacturer of the instructions, the signal was detected using the ECL system, and then the bands were captured through a CCD camera using the ChemiDoc system (Bio-Rad). Densitometric quantification of bands was done with NIH ImageJ software.

**Supplementary Figure 1**

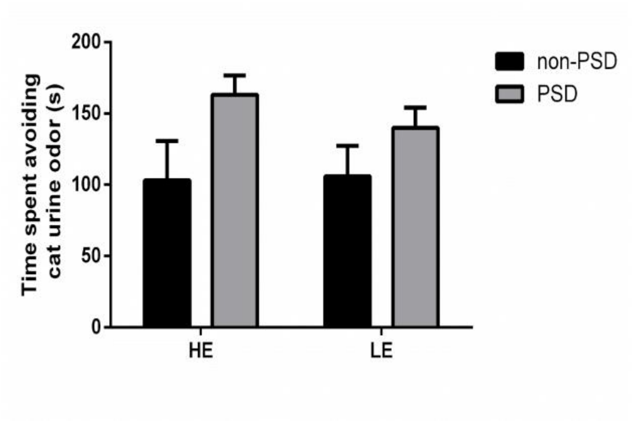

**Supplementary Table 1**

Minimum, maximum, median, mean, standard deviation, 25th, 50th and 75th percentiles of the animals' temperament division.

| | Minimum | 25% Percentile | 50% Percentile | 75% Percentile | Maximum | Mean | Std. Deviation |
|---|---|---|---|---|---|---|---|
| **HE** | 71,00 | 79,00 | 81,00 | 100,0 | 130,0 | 88,00 | 16,79 |
| **ME** | 40,00 | 43,75 | 50,50 | 60,00 | 63,00 | 51,88 | 8,509 |
| **LE** | 21,00 | 23,25 | 30,00 | 32,00 | 33,00 | 28,38 | 4,658 |

**Supplementary Table 2**

Primers used forward and reverse for q-PCR analysis.

| Gene | Forward primer | Reverse primer |
|---|---|---|
| ***Clock*** | ATGGGAATCCTTCGACACAG | GACCTTGGAAGGGTCAGTCA |
| ***Bmal 1*** | CCACAGCACAGGCTACTTGA | GCTGTGGAACCATGTGTGAG |
| ***Per 1*** | GCTGCTCCTACCAGCAAATC | AGGAGGCACATTTACGCTTG |
| ***Per 2*** | TTGTGCCTCCCGATGATGAA | AGTGGGCAGCCTTTCGATTA |
| ***Cry 1*** | GATTGACGCCATCATGACAC | CATCCCTTCTTCCCAACTGA |
| ***Cry 2*** | GGGACTACATCCGGCGATAC | ACTTAGCGGCCTTCTGAACC |
| ***Tph2*** | TGCTCAAGTTTCAGACCACCA | CCACGGCACATCCTCTAGTT |
| ***InoS*** | AGGCCACCTCGGATATCTCT | TGGGTCCTCTGGTCAAACTC |

Clock: Circadian Locomotor Output Cycles Kaput, Bmal1: Brain and Muscle ARNT-Like Protein 1, Per1: Period Circadian Regulator, Per2: Period Circadian Regulator 2, Per3: Period Circadian Regulator 3, Cry1: Cryptochrome Circadian Regulator 1 and Cry2: Cryptochrome Circadian Regulator 2, TPH2: Tryptophan Hydroxylase 2, iNOS: Inducible Nitric Oxide Synthase.

**Supplementary Table 3 – Summary of the data present in the study results section**

| Fig. # | Mean | SD | N | Statistical method used |
|---|---|---|---|---|
| **Fig. 2B- Cat odor test (Time in seconds of cloth contact)** | | | | |
| LE non-PSD | 14.3 | 11.1 | 6 | Regular 2way ANOVA Tukey's multiple comparisons test |
| LE PSD | 76 | 20.8 | 4 | |
| HE non-PSD | 66.4 | 11.1 | 5 | |
| HE PSD | 127.1 | 35.2 | 7 | |
| **Fig. 2D- Cat odor test (Time in seconds of avoidance)** | | | | |
| LE non-PSD | 105.9 | 60.5 | 8 | Regular 2way ANOVA Tukey's multiple comparisons test |
| LE PSD | 139.89 | 37.8 | 7 | |
| HE non-PSD | 103 | 73.6 | 7 | |
| HE PSD | 163 | 38.4 | 8 | |
| **Fig. 3B- Open Field test (PN30)** | | | | |
| LE | 47.5 | 9.2 | 40 | Regular 2way ANOVA Tukey's multiple comparisons test |
| ME | 80.6 | 9.9 | 20 | |
| HE | 21.9 | 7.5 | 20 | |
| **Fig. 3C- Open field test (PN60)** | | | | |
| LE | 28.4 | 4.7 | 8 | Regular 2way ANOVA Tukey's multiple comparisons test |
| ME | 51.9 | 8.5 | 8 | |
| HE | 88 | 16.8 | 8 | |

**Supplementary Table 3 continued**

| Fig. # | Mean | SD | N | Statistical method used |
|---|---|---|---|---|
| **Fig. 3D- Open Field test (Rearing)** | | | | |
| LE non-PSD | 5.8 | 1.8 | 8 | Regular 2way ANOVA Tukey's multiple comparisons test |
| LE PSD | 26.2 | 4.6 | 5 | |
| HE non-PSD | 23 | 5.6 | 9 | |
| HE PSD | 85.8 | 9.8 | 8 | |
| **Fig. 3E- Open Field test (Crossing)** | | | | |
| LE non-PSD | 56.8 | 8.4 | 5 | Regular 2way ANOVA Tukey's multiple comparisons test |
| LE PSD | 64.4 | 7.2 | 5 | |
| HE non-PSD | 90.8 | 12.8 | 6 | |
| HE PSD | 131.2 | 18.8 | 6 | |
| **Fig. 3F- Open Field test (Time spent in the central area)** | | | | |
| LE non-PSD | 31.9 | 9.9 | 6 | Regular 2way ANOVA Tukey's multiple comparisons test |
| LE PSD | 30.3 | 12.4 | 5 | |
| HE non-PSD | 34.1 | 8.6 | 5 | |
| HE PSD | 51.7 | 8.9 | 6 | |
| **Fig. 3G- Open Field test (Ratio time spent in the central area/total time spent in the open field)** | | | | |
| LE non-PSD | 0.2 | 0.1 | 6 | Regular 2way ANOVA Tukey's multiple comparisons test |
| LE PSD | 0.2 | 0.01 | 5 | |
| HE non-PSD | 0.2 | 0.01 | 6 | |
| HE PSD | 0.4 | 0.1 | 5 | |
| **Fig. 3H- Open Field test (Freezing)** | | | | |
| LE non-PSD | 11.3 | 3.3 | 6 | Regular 2way ANOVA Tukey's multiple comparisons test |
| LE PSD | 10.6 | 4.0 | 4 | |
| HE non-PSD | 9.6 | 1.9 | 6 | |
| HE PSD | 10.8 | 2.5 | 6 | |

**Supplementary Table 3 continued**

| Fig. # | Mean | SD | N | Statistical method used |
|---|---|---|---|---|
| **Fig. 4B- Elevated Plus-maze test (Number of entries into the open arms)** | | | | |
| LE non-PSD | 120 | 63.2 | 10 | Regular 2way ANOVA Tukey's multiple comparisons test |
| LE PSD | 200 | 57.7 | 7 | |
| HE non-PSD | 290 | 87.6 | 10 | |
| HE PSD | 557.1 | 151.2 | 7 | |
| **Fig. 4C- Elevated Plus-maze test (Time spent in the open arms %)** | | | | |
| LE non-PSD | 37.2 | 15.2 | 9 | Regular 2way ANOVA Tukey's multiple comparisons test |
| LE PSD | 46.1 | 5.2 | 8 | |
| HE non-PSD | 54.1 | 6.5 | 9 | |
| HE PSD | 87.7 | 13.9 | 8 | |
| **Fig. 4D- Elevated Plus-maze test (Number of entries into the closed arms)** | | | | |
| LE non-PSD | 220.0 | 122.98 | 10 | Regular 2way ANOVA Tukey's multiple comparisons test |
| LE PSD | 337.5 | 140.8 | 8 | |
| HE non-PSD | 460.0 | 117.4 | 10 | |
| HE PSD | 562.5 | 140.8 | 8 | |
| **Fig. 5B- Y maze test (incorrect alternations %)** | | | | |
| LE non-PSD | 41.9 | 6.6 | 8 | Regular 2way ANOVA Tukey's multiple comparisons test |
| LE PSD | 76.1 | 4.2 | 8 | |
| HE non-PSD | 54.1 | 6.5 | 9 | |
| HE PSD | 76.6 | 4.2 | 6 | |
| **Fig. 5D- Novel object recognition (Discrimination ratio)** | | | | |
| LE non-PSD | 0.7 | 0.1 | 6 | Regular 2way ANOVA Tukey's multiple comparisons test |
| LE PSD | -0.6 | 0.5 | 5 | |
| HE non-PSD | 0.9 | 0.08 | 6 | |
| HE PSD | 0.04 | 0.06 | 6 | |

**Supplementary Table 3 continued**

| Fig. # | Mean | SD | N | Statistical method used |
|---|---|---|---|---|
| **Dopamine receptor 1** | | | | |
| LE non-PSD | 1.1 | 0.3 | 3 | Bonferroni's multiple comparisons test |
| LE PSD | 0.7 | 0.1 | 3 | |
| HE non-PSD | 1.0 | 0.2 | 3 | |
| HE PSD | 2.6 | 0.8 | 4 | |
| **Dopamine receptor 2** | | | | |
| LE non-PSD | 1.4 | 0.4 | 3 | Bonferroni's multiple comparisons test |
| LE PSD | 2.5 | 0.6 | 4 | |
| HE non-PSD | 1.3 | 0.7 | 4 | |
| HE PSD | 1.2 | 0.3 | 4 | |

**Figure S3 – Full-length gels for D1 and D2**

D1

Membrane 1

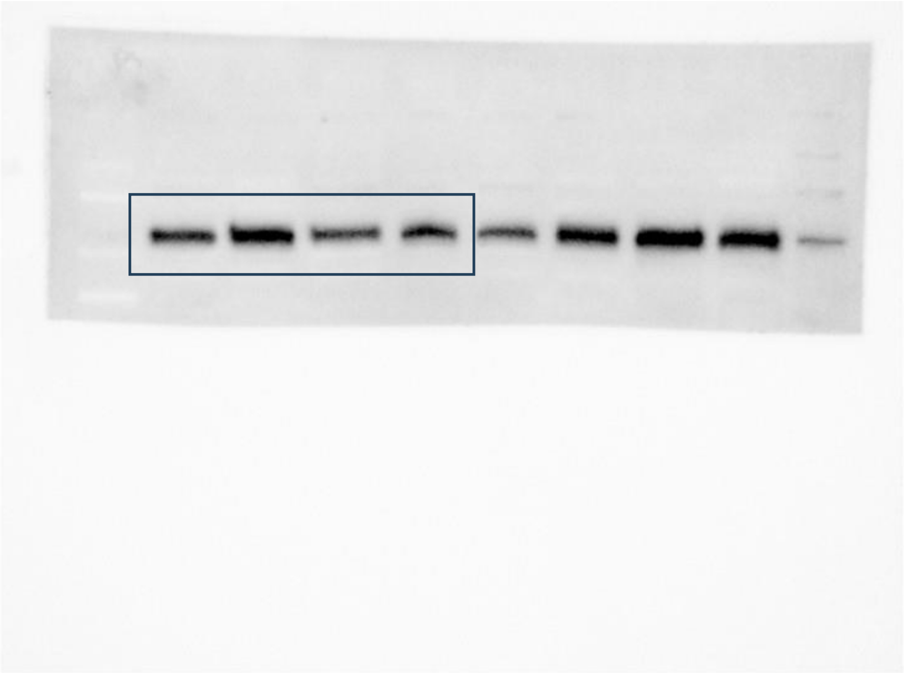

Membrane 2

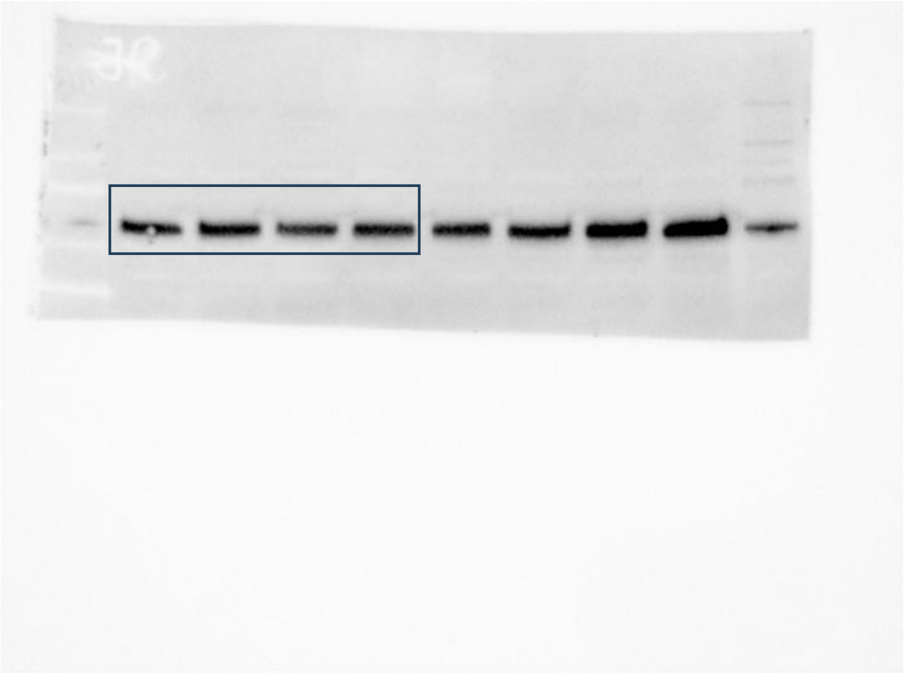

Membrane 3

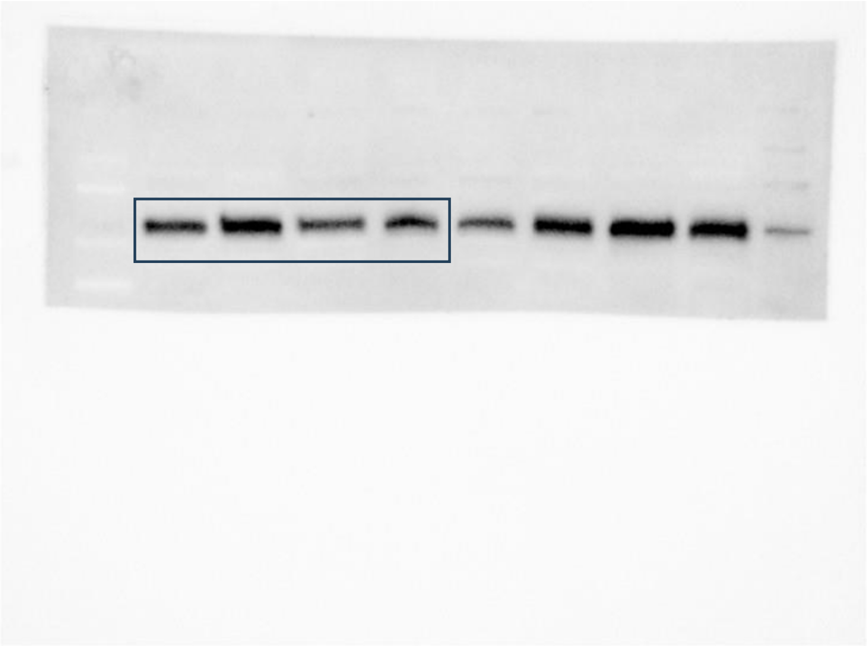

Membrane 4

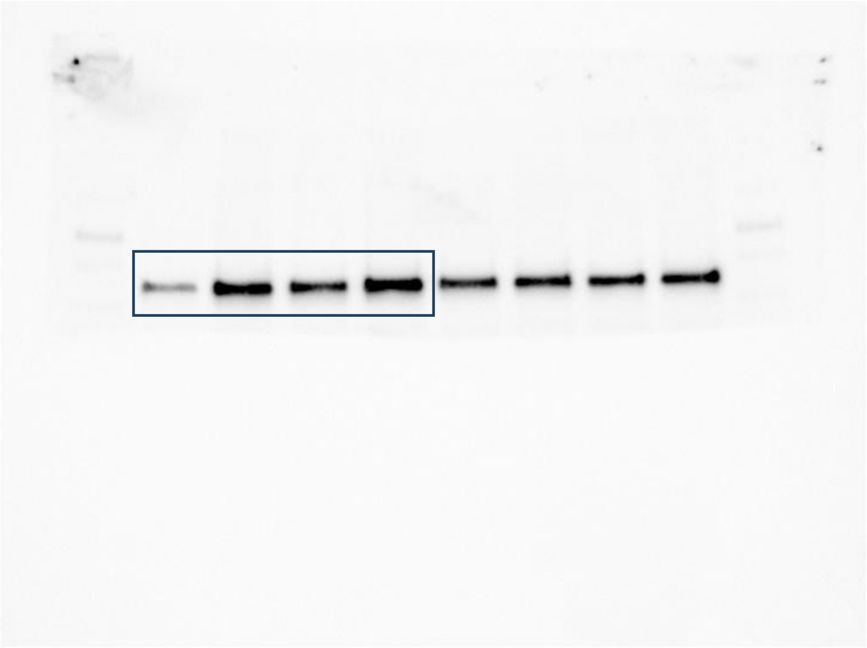

D2

Membranes 1 and 2

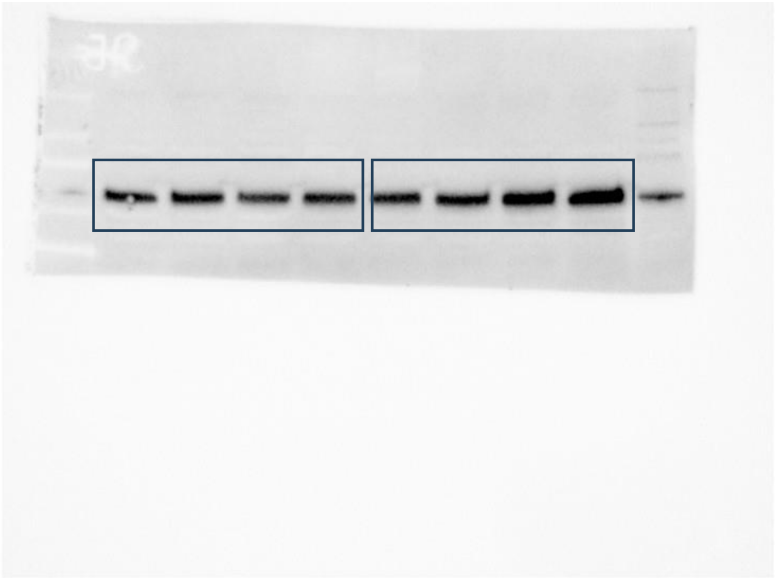

Membrane 3

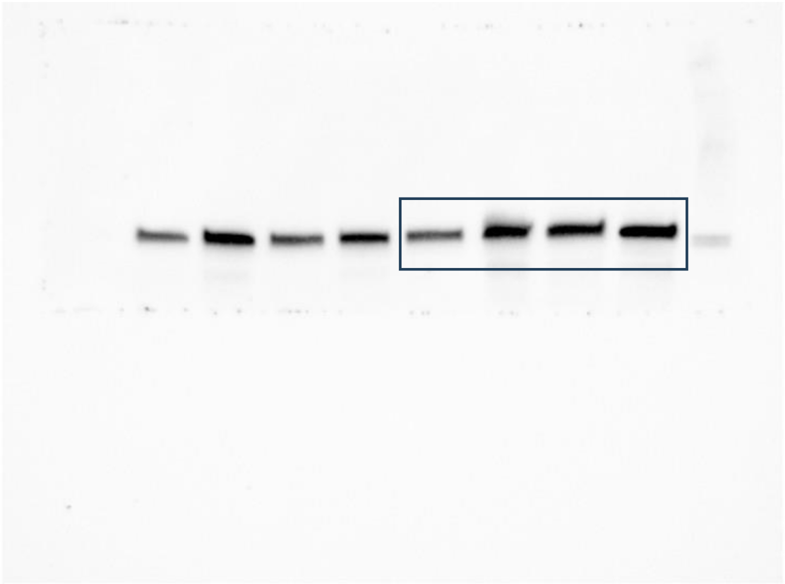

Membrane 4

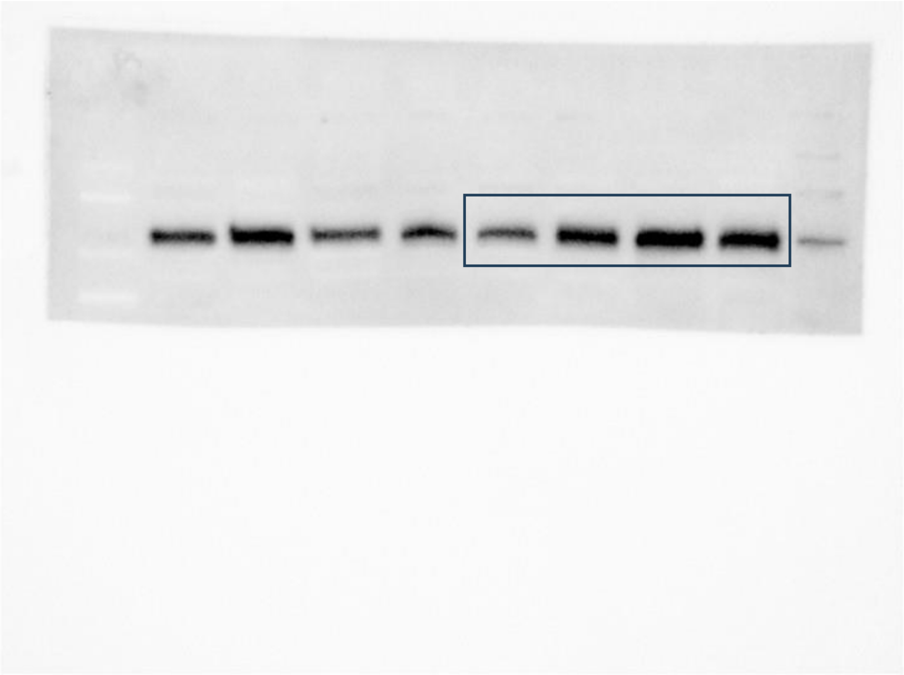
